# Bidirectional Upregulation of Klotho by Triiodothyronine and Baicalein: Mitigating Chronic Kidney Disease and Associated Complications in Aged BALB/c Mice

**DOI:** 10.1101/2024.09.08.611868

**Authors:** Saswat Kumar Mohanty, Vikas Kumar Sahu, Bhanu Pratap Singh, Kitlangki Suchiang

## Abstract

Chronic kidney disease (CKD) is a global health challenge marked by progressive renal decline and increased mortality. The interplay between CKD and hypothyroidism, particularly nonthyroidal low-triiodothyronine (T3) syndrome, exacerbates disease progression, driven by HPT axis dysfunction and reduced Klotho levels due to Wnt–β-catenin pathway activation. This study explored Klotho as a link between CKD and hypothyroidism using an adenine-induced CKD aged mouse model. Exogenous T3 and baicalein (BAI), targeting the Wnt pathway, were used to upregulate Klotho expression. Combined T3 and BAI treatment significantly increased Klotho levels, surpassing individual effects, and suppressed key signaling molecules (TGF, NFκB, GSK3), mitigating renal fibrosis and CKD complications, including cardiovascular disorders and dyslipidemia. This bidirectional approach, enhancing Klotho via T3 and sustained Wnt pathway inhibition, offers a novel and effective strategy for CKD management, particularly in elderly patients with hypothyroidism.

## Introduction

The prevalence of chronic kidney disease (CKD) has accelerated worldwide and is becoming a severe health problem [1]. Recent studies suggest that an abnormal increase in CKD, especially in developing Asian countries, is associated with hypertension, type 2 diabetes, and cardiovascular disease (CVD) [2]. The increasing CKD trend can be correlated with the aging population in the last several decades. Consistent with this concept, many studies suggest that aging is an independent risk factor for the development and progression of CKD [3,4]. Many reports also indicate that CKD is associated with numerous complications, including dyslipidemia, CVD, mineral bone disorder, and thyroid dysfunction [5]. The kidney plays an essential role in thyroid hormone metabolism, degradation, and excretion [6]. Reduced expression of both triiodothyronine (T3) and thyroxine (T4) with elevated levels of thyroid-stimulating hormone (TSH) is commonly observed in CKD patients because of dysregulation of the hypothalamus□pituitary□thyroid (HPT) axis [7].

Klotho is a single-pass transmembrane protein detectable in limited tissues, particularly in distal convoluted tubules of the kidneys and the choroid plexus in the brain. The Klotho protein may regulate the occurrence of age-related disorders or natural aging processes by functioning as a circulating humoral factor [8]. The Klotho protein is approximately 130 kDa (1,014 amino acids) with a presumed signal sequence at the N-terminus, a single transmembrane domain at the C-terminus, and a short cytoplasmic domain (10 amino acids) [9,10]. Membrane Klotho is proteolytically cleaved and shed by different types of secretases, such as α, β, and γ secretases [11–13]. There is also a small secreted form of Klotho produced by alternative splicing consisting of 550 amino acids, a truncated Klotho protein [14]. Clinical studies have suggested that both the membrane and secreted forms of Klotho are downregulated in human CKD, demonstrating that the loss of Klotho is correlated with the pathogenesis of kidney injury [15,16].

Building upon this foundation, recent investigations have sought to elucidate the mechanistic link between CKD and hypothyroidism through the lens of Klotho biology. Notably, our prior studies revealed that T3, a thyroid hormone, could increase Klotho expression in *Caenorhabditis elegans* (*C. elegans*).[17] This raises intriguing questions about the potential benefits of T3 supplementation in mitigating the impaired HPT axis and conferring protective effects against age-associated CKD, particularly in higher mammalian models.

Moreover, many studies reported the over activation of the Wnt/β-catenin pathway in kidney disease including CKD. This overactivation also leads to kidny fibrosis by diposition of ESM. [18–23]. Overactivation of the Wnt pathway in kidney diseases also downregulates Klotho expression [24]. In contrast, Klotho can protect against CKD by inhibiting the Wnt pathway [25–27].

Furthermore, in our prior study involving the nematode *C. elegans*, we discovered a significant association: suppression of the Wnt pathway leads to increased Klotho expression [28]. Building on these observations, our previous research revealed that T3 can notably augment various forms of kidney-specific Klotho protein while reducing the increase in Wnt/β-catenin pathway activity in aged mice [29]. This reciprocal regulatory mechanism prompts inquiries into its evolutionary conservation across species and potential implications for chronic kidney disease. To supress the inhibition of Klotho by the Wnt/β-catenin pathway in CKD, we chosed a natural compound baicalein (BAI) which is known to inhibits the Wnt/β-catenin pathway. [30–33]. Moreover, preliminary studies from our laboratory using the nematode *C. elegans* revealed that BAI could increase Klotho expression by inhibiting Wnt ligands [28].

Hence, we assume that stabilizing Klotho expression in CKD kidney may be benificial and can be used as a theraputic approach for CKD patients.So we proposes a novel bidirectional targeting approach that leverages both exogenous hormone (T3) and the natural compound BAI to achieve stable upregulation of Klotho. The rationale behind this dual intervention is grounded in the potential synergistic effects of hormonal and natural compound interventions, aiming for enhanced protective effects against CKD.

## Materials and methods

### Animal care and treatment

Animal studies were performed after approval by the Institutional Animal Ethics Committee (IAEC) of Pondicherry University, India (IAEC approval ref. No: PU/CAHF/23rd IAEC/2019/03, dated 15.04.2019). All the experiments were conducted in accordance with the guidelines of the Committee for the Control and Supervision of Experiments on Animals (CPCSEA), New Delhi, India, under the supervision of the IAEC. The animals were housed under conventional conditions: 23^0^ ± 2^0^ C, 12:12 light:dark cycle, standard diet, and water *ad libitum*. Twenty-month-old BALB/c mice were used for this study. After 15 days of acclimatization, the animals were grouped, with six animals in each group, on the basis of their weight. Blood collection and euthanasia of the animals were carried out under isoflurane narcosis. All experimental animals were approved in advance by the Institutional Animal Ethics Committee (IAEC). CKD was induced by the addition of 50 mg/kg b.w. adenine (Sigma Aldrich, USA) dissolved in 0.5% carboxy methyl cellulose (CMC) via an oral dose for 21 days, whereas the control animals received 0.5% CMC for only 21 days Add reference [34]. After CKD induction, the animals were further divided into different groups. The CKD group received a vehicle for T3 and BAI (0.1 N NaOH and DMSO) and was named the placebo group. The other groups received an individual dose of either 1 mg/kg b.w. T3 or 75 mg/kg b.w. Baicalein (BAI) (Sigma Aldrich, USA) was administered after treatment. The T3 and BAI doses were fixed based on previous reports [35–39]. However, the supraphysiologic dose of T3 has been chosen to monitor its physiological effect on Klotho protein expression in the kidney. The last group received the combined treatment (T3 + BAI group). BAI was administered for five consecutive days after 21 days, whereas T3 was administered during the last three days, i.e., the 24th–26th days. All the animal experiments used in the manuscript followed the recommendations of the ARRIVE guidelines. The experimental design is outlined in **Fig. 1**.

**Fig. 1.**
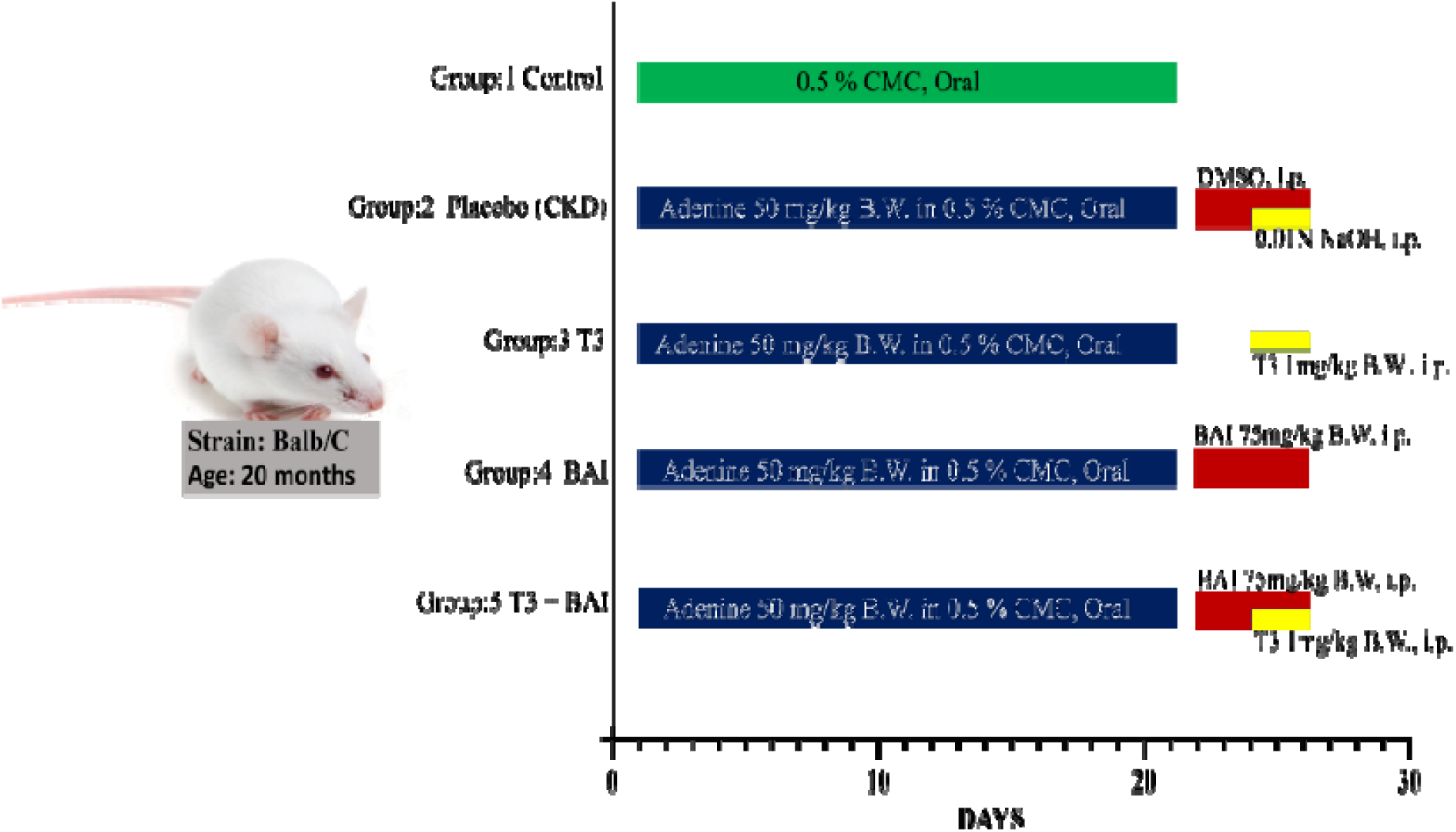
Animal grouping and treatments.

### Serum and urine biochemical parameters

Serum creatinine, urea, blood urea nitrogen (BUN), albumin, calcium, inorganic phosphorus, hemoglobin, total cholesterol (TC), total triglyceride (TG), high-density lipoprotein (HDL-C), alkaline phosphatase (ALP) activity, bilirubin, urine creatinine, and urea were measured via the Agappe diagnostic kit (Agappe Diagnostic, India) according to the manufacturer’s instructions. LDL-C and VLDL-C were calculated via the equation provided by Sampson et al. [40] as follows:

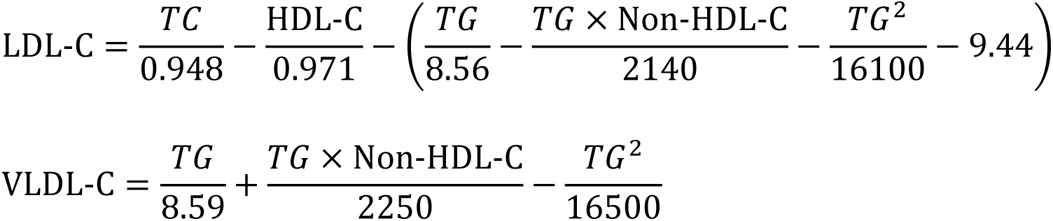

### ELISA

T3 (E-EL-0079), TSH (E-EL-M1153), and 1,25 hydroxy Vit-D3 (E-EL-0016) ELISA kits were purchased from Elabscience USA. T3 and Vit-D3 levels were measured via competitive ELISA, and TSH levels were measured via the sandwich ELISA method according to the manufacturer’s instructions.

### Histological analysis

Kidney tissue was harvested and fixed in 10% neutral buffered formalin, and a 4-µm-thick section was prepared from paraffin-embedded tissue with the help of an expert histopathologist. The kidney sections were further stained with H&E and Masson’s trichrome. Kidney damage was evaluated on the basis of parameters such as inflammatory infiltration, protein casts, interstitial fibrosis, and ECM deposition. The parameters were measured in ten randomly selected fields from each section at 400x magnification. All the parameters were measured via ImageJ software.

### RNA isolation and qRT□PCR analysis

RNA express reagent (Hi-Media, India) was used to isolate total RNA from mouse kidney tissue. Briefly, 2 µg of RNA was subjected to cDNA synthesis via a cDNA isolation kit (Thermo Fisher Scientific, USA) per the manufacturer’s instructions. cDNA was subjected to qPCR using SYBR green master mix (Roche, USA). The qPCR was run on a Hi-Media Insta Q48^TM^ real-time machine for 40 cycles. mRNA was quantified via the 2^-ΔΔ*Ct*^ method, with β-actin used as an internal control [41]. The details of the primers used are listed in **Table 1**.

**Table 1.**
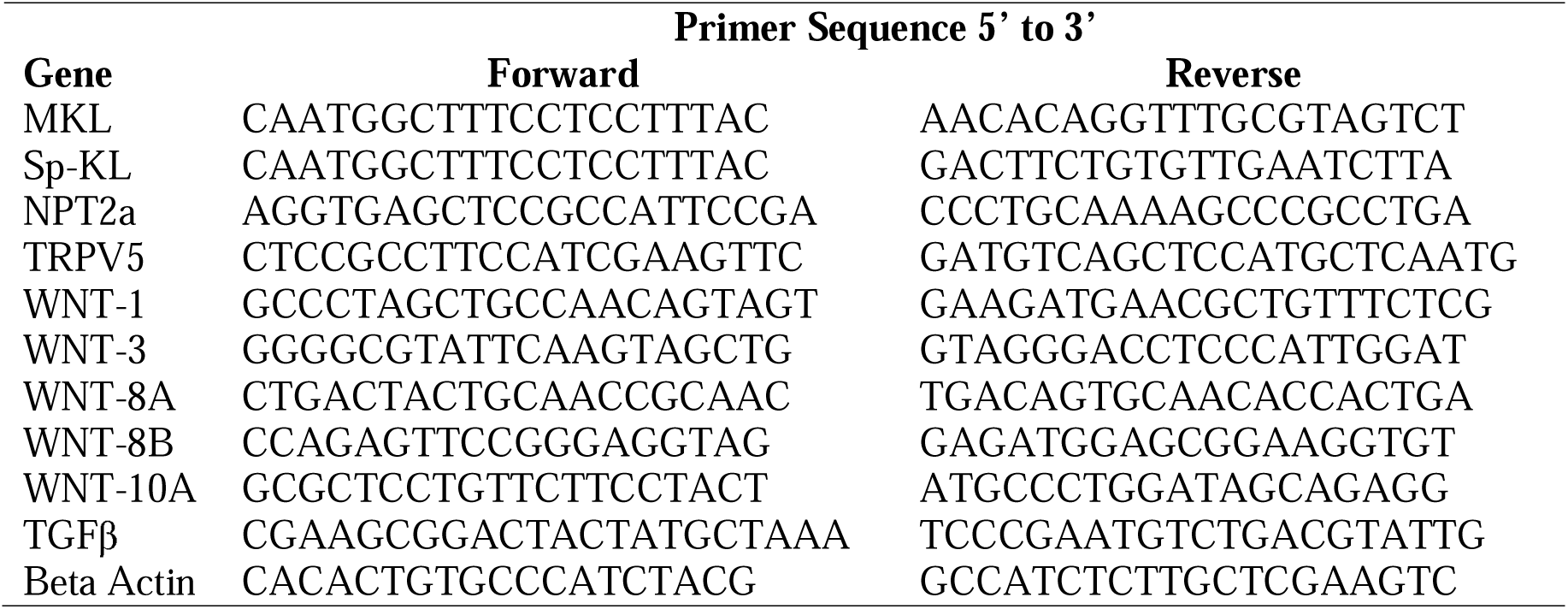
Mouse nucleotide sequences of the primers used for qRT□PCR analysis.

### Western blotting analysis

For western blot analysis, all reagents were purchased from Sigma Aldrich, USA, unless otherwise indicated. Kidney tissue was homogenized in RIPA lysis buffer supplemented with protease and phosphatase inhibitors. Equal concentrations of proteins were loaded and electrophoresed on a 10% gradient tris-glycine gel. Proteins were electrophoretically transferred to an immobile-P PVDF membrane (Roche, USA). After being blocked with 5% nonfat skim milk for 1 h at room temperature, the membranes were incubated with primary antibodies (Santacruz, USA) against anti-Klotho (sc-515942), anti-GSK3β (sc-7291), anti-β-catenin (sc-7963), anti-CK1 (sc-74582), anti-NFκB (sc-8008), anti-IL6 (sc-57315) and anti-GAPDH (sc-32233) at 4°C overnight. After thorough washing with buffers, the membranes were incubated with a secondary antibody (sc-516102) for 1 h at room temperature. The bands were then developed in a chemiluminescence substrate via the Vilber Chemi Doc imaging system. The intensity of each band was measured via ImageJ software, and the relative expression was determined with respect to that of GAPDH.

### Antioxidant enzyme assay

#### Catalase assay

Catalase activity was measured as described previously [42]. The tissue homogenate was incubated in the substrate at 37°C for three minutes. After incubation, the reaction was stopped with ammonium molybdate. The absorbance of the molybdate and hydrogen peroxide yellow complex was measured at 374 nm against a blank. The specific activity was expressed as U/mg of protein.

#### SOD activity

SOD activity in the tissue homogenate was measured via the methods of Kostyuk V A et al. [43]. SOD activity was measured by the inhibition of autooxidation of quercetin by the enzyme at pH 10. The results are expressed as U/mg of protein.

### Statistical analysis

Two-way analysis of variance with post hoc Tukey correction was used to compare the means of different groups. All experiments were performed in triplicate, and the results are reported as the means□±□SEMs. The graphs were plotted via GraphPad Prism software (GraphPad Software, San Diego, CA, USA), and statistical significance was also measured via the same software. The error bars represent the SEM. p values < 0.05 were considered statistically significant.

## Results

### CKD reduces body weight and improves with bidirectional treatment

The body weights of all the animals were measured every two days during the treatment period. The body weights of the animals significantly decreased during the adenine administration period. At the adenine diet endpoint (21st day), the adenine-fed placebo group animals weighed 31% less than the control group did (p<0.001). In contrast, when adenine-fed animals were treated with T3, BAI, or the combined dose (T3 + BAI) alone, significant improvements in body weight at the end of the treatments (27th day) were observed. The T3-treated animals presented an increase in body weight of 17.9% (p<0.001), the BAI-treated animals presented an increase of 16.6% (p<0.001), and the combined (T3 + BAI)-treated animals presented the greatest increase of 19.2% (p<0.001) from the 27th-day placebo group body weight **(Fig. 2).**

**Fig. 2.**
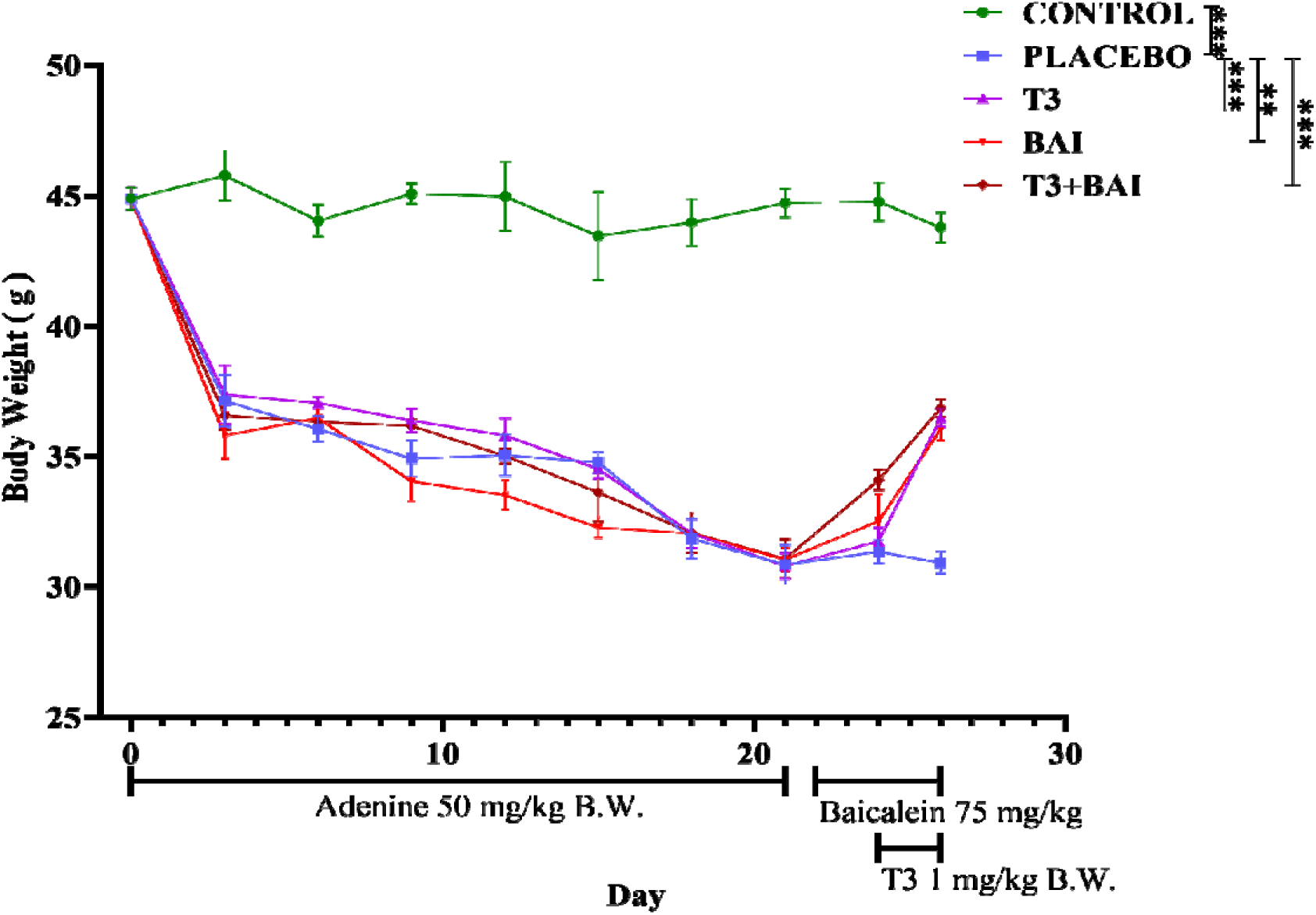
Body weight profile of BALB/c mice during CKD induction and treatment. Body weight was measured every 2 days during the treatment. A significant reduction in body weight was observed after adenine treatment. However, treatment with either T3, baicalein (BAI) or the combination (T3 + BAI) significantly increased the body weights of the animals. n = 10 per group. ⍰⍰ and ⍰⍰⍰ indicate significance at p ≤ 0.01 and p ≤ 0.001, respectively.

### Bidirectional treatment ameliorates abnormal serum and urine parameters in CKD animals

The present study of adenine-induced CKD in aged animals assessed kidney injury via serum and biochemical kidney injury markers. We initially observed that the serum creatinine, urea, and BUN concentrations were significantly greater in the placebo group than in the control group. The percentage of increase in creatinine was 8.8-fold (p<0.001), that in urea was 1.56-fold (p<0.05), and that in BUN was 1.56-fold (p<0.05). In contrast, the urine creatinine and urea concentrations in the placebo group were significantly lower than those in the control group (3.12-fold (p<0.01) and 2.08-fold (p<0.01), respectively). Moreover, the urine ALB concentration increased 3.05-fold (p<0.001) in the placebo group than in the control group, leading to significant albuminuria. However, treatment with either T3, BAI or their combination (T3 + BAI) ameliorated the changes observed in the placebo group toward the normal range. Specifically, in comparison with the treated groups, the combined (T3 + BAI) treatment produced a more pronounced effect, in which the serum creatinine, urea and BUN levels were 3.8-fold (p<0.01), 1.63-fold, (p<0.01), and 1.68-fold (p<0.001) lower, respectively, than those in the placebo group. **(Table 2)**

**Table 2.**
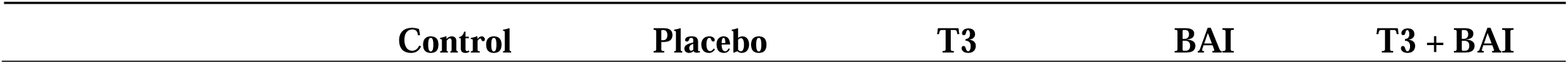
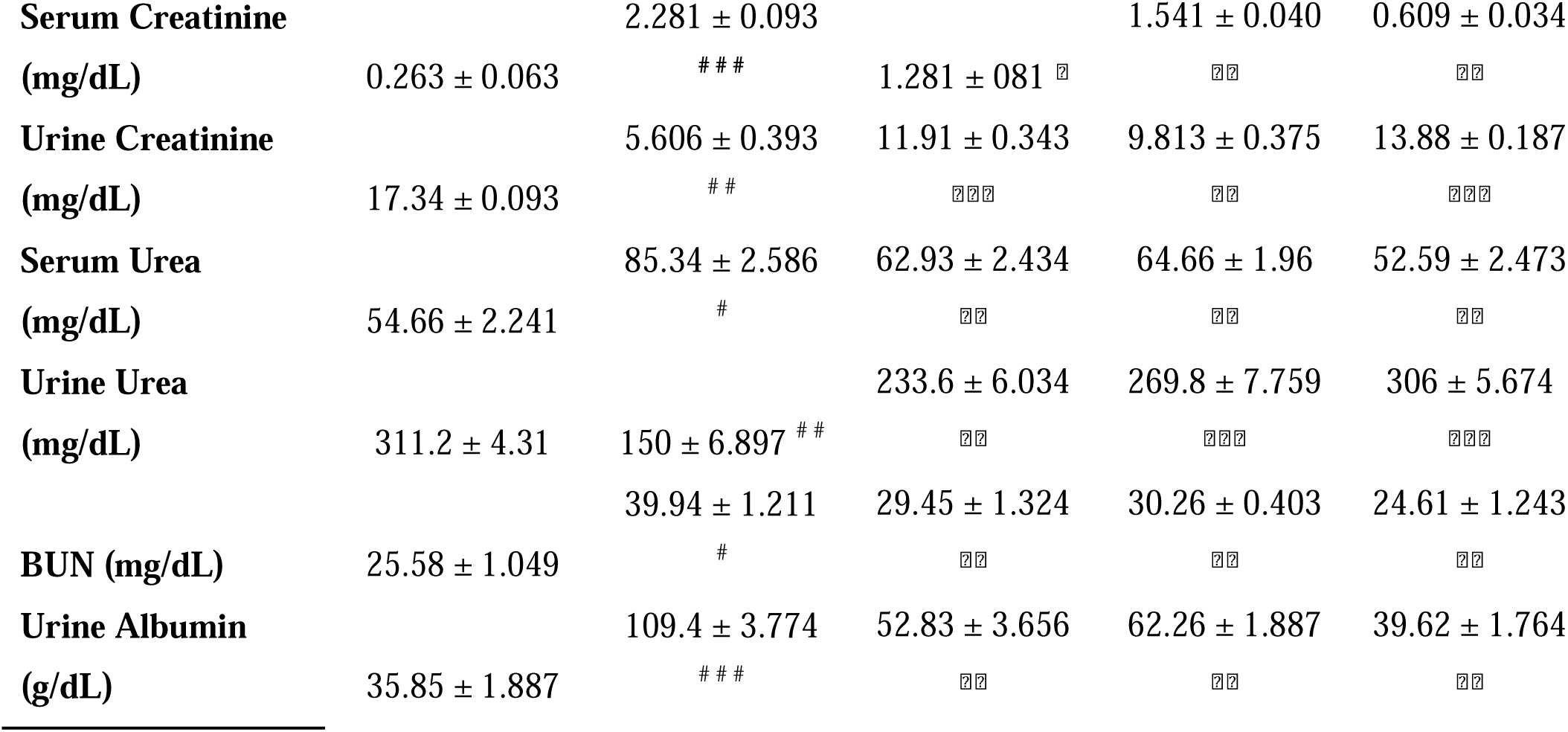
Kidney function test: Serum and urine biochemical parameters.

| Control | Placebo | T3 | BAI | T3 + BAI |
| --- | --- | --- | --- | --- |

|  |  |  |  |  |  |
| --- | --- | --- | --- | --- | --- |
| **Serum Creatinine** |  | 2.281 ± 0.093 |  | 1.541 ± 0.040 | 0.609 ± 0.034 |
| **(mg/dL)** | 0.263 ± 0.063 | ### | 1.281 ± 0.081 |  |  |
| **Urine Creatinine** |  | 5.606 ± 0.393 | 11.91 ± 0.343 | 9.813 ± 0.375 | 13.88 ± 0.187 |
| **(mg/dL)** | 17.34 ± 0.093 | ## |  |  |  |
| **Serum Urea** |  | 85.34 ± 2.586 | 62.93 ± 2.434 | 64.66 ± 1.96 | 52.59 ± 2.473 |
| **(mg/dL)** | 54.66 ± 2.241 | # |  |  |  |
| **Urine Urea** |  |  | 233.6 ± 6.034 | 269.8 ± 7.759 | 306 ± 5.674 |
| **(mg/dL)** | 311.2 ± 4.31 | 150 ± 6.897 |  |  |  |
|  |  | 39.94 ± 1.211 | 29.45 ± 1.324 | 30.26 ± 0.403 | 24.61 ± 1.243 |
| **BUN (mg/dL)** | 25.58 ± 1.049 | # |  |  |  |
| **Urine Albumin** |  | 109.4 ± 3.774 | 52.83 ± 3.656 | 62.26 ± 1.887 | 39.62 ± 1.764 |
| **(g/dL)** | 35.85 ± 1.887 | ### |  |  |  |

### Effects of T3, BAI, and the combined treatment (T3 + BAI) on kidney fibrosis and extracellular matrix deposition in CKD animals

Kidney sections were subjected to H&E and Masson’s trichrome staining to assess pathological renal conditions **(Fig. 3).** H&E staining **(Fig. 3a)** revealed that CKD mice in the placebo group presented significant monocyte and lymphocyte infiltration (A), protein casts (B), and cytoplasmic vacuolation (C) in renal sections. Moreover, the Masson’s trichrome-stained **(Fig. 3b)** renal sections indicate an increase in interstitial fibrosis with a high deposition of extracellular matrix components in the placebo group. However, treatment with T3, BAI or their combination (T3 + BAI) significantly improved the damage. The damage to the renal sections was evaluated on the basis of ten randomly selected fields per section, which were examined at 400x magnification. The evaluation criteria and observed results are presented in **Table 3**. Overall, our results showed that the combined treatment produced a better counteractive effect on renal fibrosis and ECM accumulation.

**Fig. 3.**
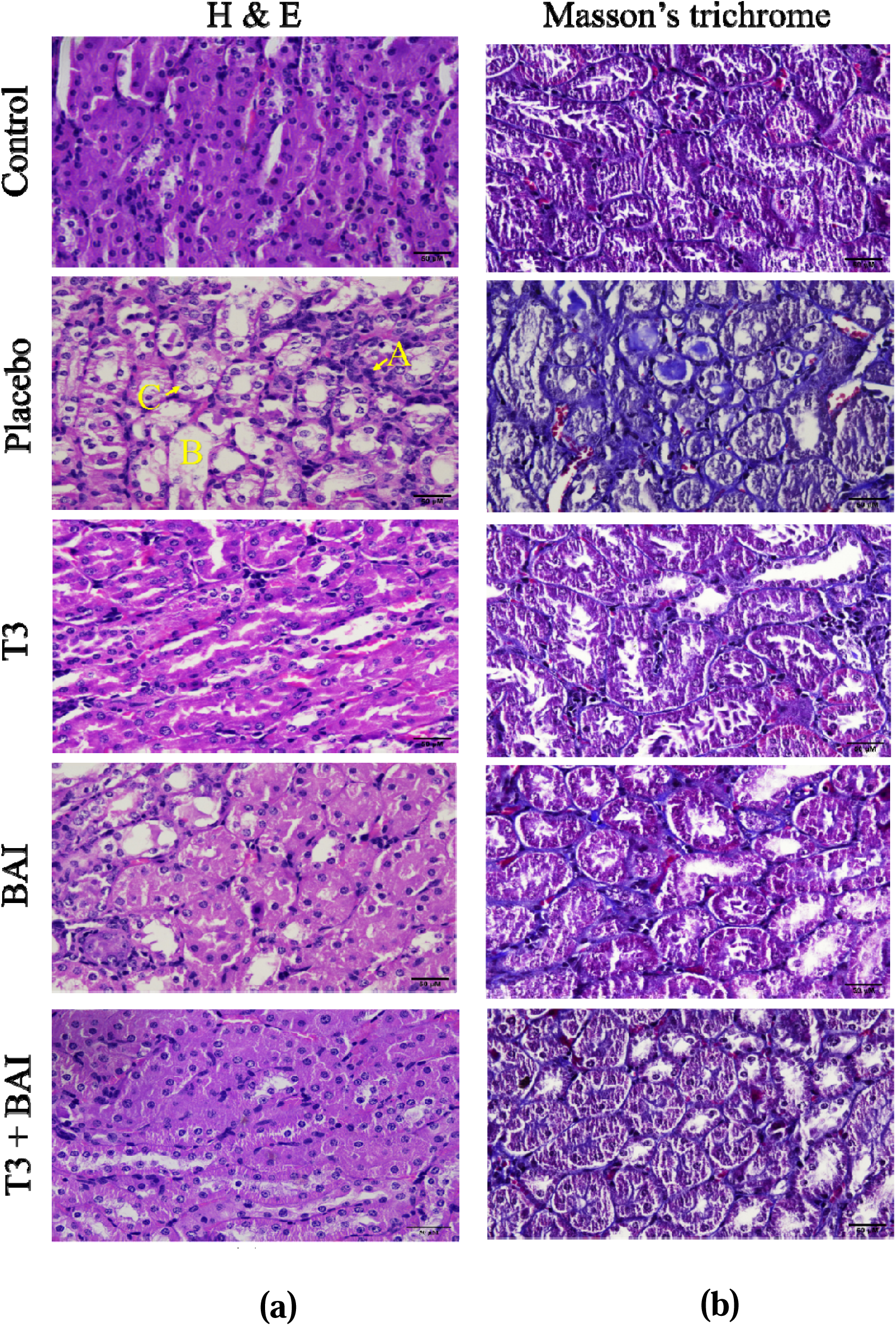
Analysis of renal pathological sections from CKD patients. These findings indicate the protective effects of T3, BAI, and their combination (T3 + BAI) against CKD-induced renal fibrosis. **(a)** Hematoxylin and eosin (H&E) staining of renal pathological sections (x400), **(b)** Masson’s trichrome staining for extracellular matrix deposition analysis (x400). **A, B** and **C** represent inflammatory infiltration, protein casting, and cytoplasmic vacuolation, respectively. The scale bar represents 50 µM.

**Table 3.** Evaluation of the degree of damage in renal histological sections.

| Type | Criteria |  |  |  | Results |  |  |  |  |
| --- | --- | --- | --- | --- | --- | --- | --- | --- | --- |
|  | 0 | 1 | 2 | 3 | Control | Placebo | T3 | BAI | T3 + BAI |
| **Inflammatory Infiltration** | None | Scatter | Focal | Diffuse | 0 | 3 | 2 | 2 | 1 |
| **Protein Cast** | None | < 10% | 10-25% | > 25% | 0 | 2 | 1 | 1 | 1 |
| **Interstitial Fibrosis** | None | < 25% | 25-50% | > 50% | 0 | 3 | 2 | 2 | 1 |
| **ECM Deposition** | None | < 25% | 25-50% | > 50% | 0 | 3 | 1 | 2 | 1 |
| **Total** |  |  |  |  | **0** | **11** | **6** | **7** | **4** |

Each sample was performed based on ten randomly selected fields per section under x400 magnification.
The values are mean ± SEM. n = 5. □, □□, and □□□ indicate the significance level $p \leq 0.05$ , $p \leq 0.01$ and $p \leq 0.001$ , respectively from placebo. #, ##, and ### indicate the significance level $p \leq 0.05$ , $p \leq 0.01$ and $p \leq 0.001$ , respectively from control.

### Combined treatment (T3 + BAI) maximally counteracts the decline in kidney Klotho levels during CKD

Recent evidence has shown that Klotho can serve not only as an early biomarker of CKD but also as a potential therapeutic target for CKD [44,45]. In our study of adenine-induced CKD, we found that the mRNA expression of M-Klotho and Sp-Klotho was significantly lower in the placebo group than in the control group by 3.25-fold (p<0.001) and 2.35-fold (p<0.001), respectively **(Fig. 4a, c).** In contrast, M-Klotho expression increased significantly in the T3, BAI, and combined (T3 + BAI) groups by 4.5-fold (p<0.001), 3.6-fold (p<0.01), and 5.5-fold (p<0.001), respectively, compared with that in the placebo group **(Fig. 4b).** Similarly, Sp-Klotho expression also increased significantly upon T3, BAI, and combined (T3 + BAI) treatment by 4.2-fold (p<0.001), 2.4-fold (p<0.05), and 5.3-fold (p<0.001), respectively **(Fig. 4d).** The protein expression of Klotho revealed that α-KL and SKL expression was markedly lower in the placebo-treated group than in the control group (91.6% and 15.5%, respectively) **(Fig. 4e, f, g)**. Upon treatment with T3 and BAI, a significant increase in Klotho protein expression was observed, and the combined treatment (T3 + BAI) resulted in greater Klotho protein expression than either of the individual treatments did.

**Fig. 4.**
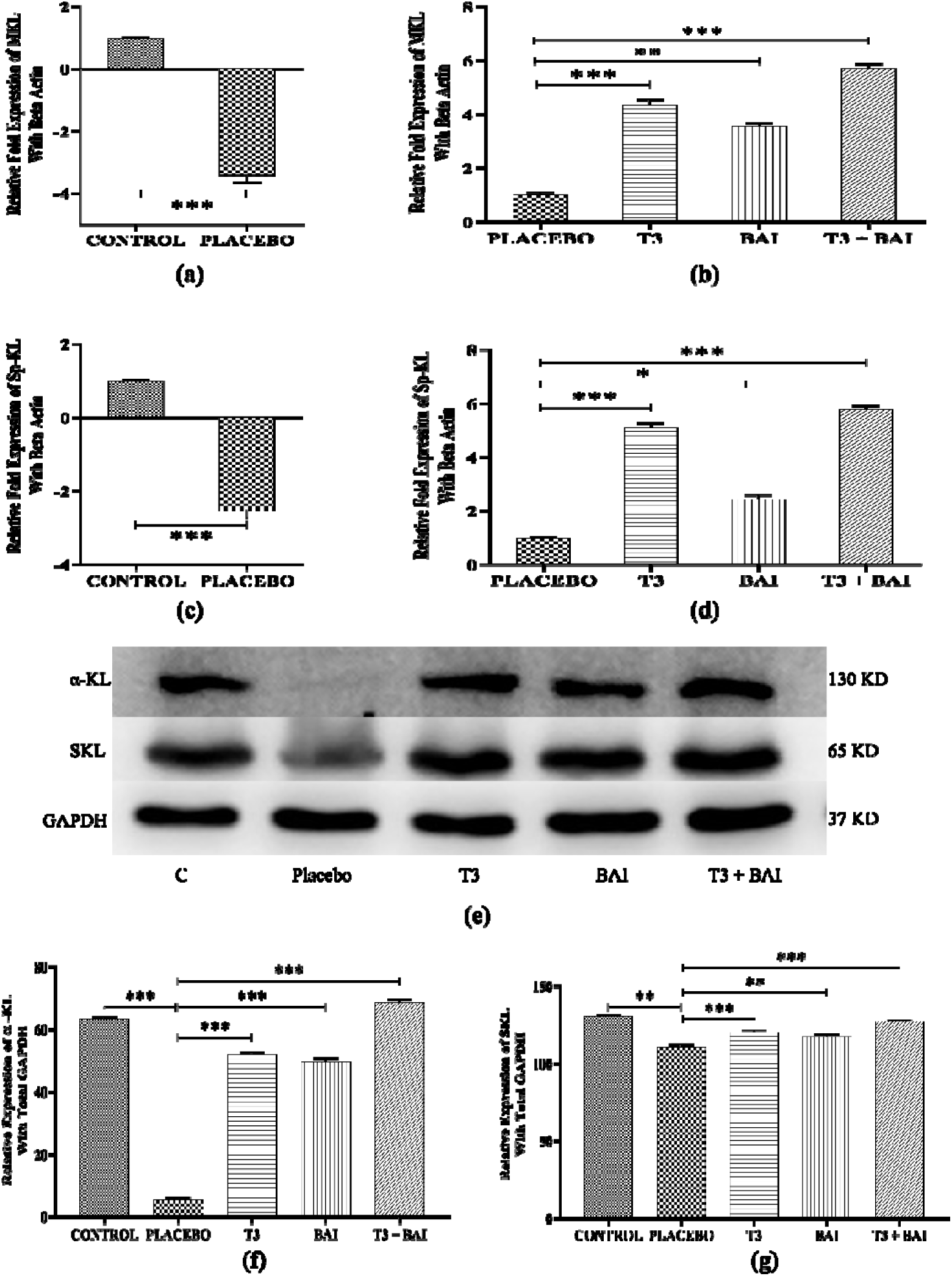
Effects of T3, BAI, and the combined treatment (T3 + BAI) on Klotho expression in CKD kidneys. Klotho mRNA, i.e., membrane Klotho (M-KL) and spliced Klotho (Sp-KL), and protein, i.e., α-KL and S-KL, expression are decreased in CKD. After treatment with T3 and BAI in CKD mice, both the mRNA and protein expression levels increased significantly, with the combined treatment (T3 + BAI) resulting in the greatest increase in expression. **(a)** and **(b)** represent M-KL mRNA expression; **(c)** and **(d)** represent Sp-KL mRNA expression. **(e)** Western blot analysis of α-KL and soluble Klotho (S-KL); **(f)** and **(g)** Densitometric analysis of α-KL and S-KL, respectively. The data are expressed as the means□±□SEMs from 3 independent experiments. ⍰, ⍰⍰, and ⍰⍰⍰ indicate significance at p ≤ 0.05, p ≤ 0.01, and p ≤ 0.001, respectively.

### Aberrant increase in GSK-3**β** expression during CKD is maximally suppressed by T3 + BAI

Recent studies have also implicated GSK3β in diverse kidney diseases, and targeting GSK3β improved kidney fibrosis [46,47]. To address this issue, we evaluated the expression of GSK-3β, which tends to be overexpressed in different types of kidney injury. We found a significant 3.3-fold (p<0.001) increase in GSK-3β protein expression in the kidneys of the adenine-fed placebo group compared with the control group. Interestingly, compared with the placebo, the T3 and BAI treatments counteracted the observed changes in expression by 2.1-fold (p<0.001) and 1.8-fold (p<0.01), respectively. The optimal downregulation of GSK3β, with a decrease of 4.6-fold (p<0.001), was observed in the combined (T3 + BAI)-treated group **(Fig. 5a, b)**.

**Fig. 5.**
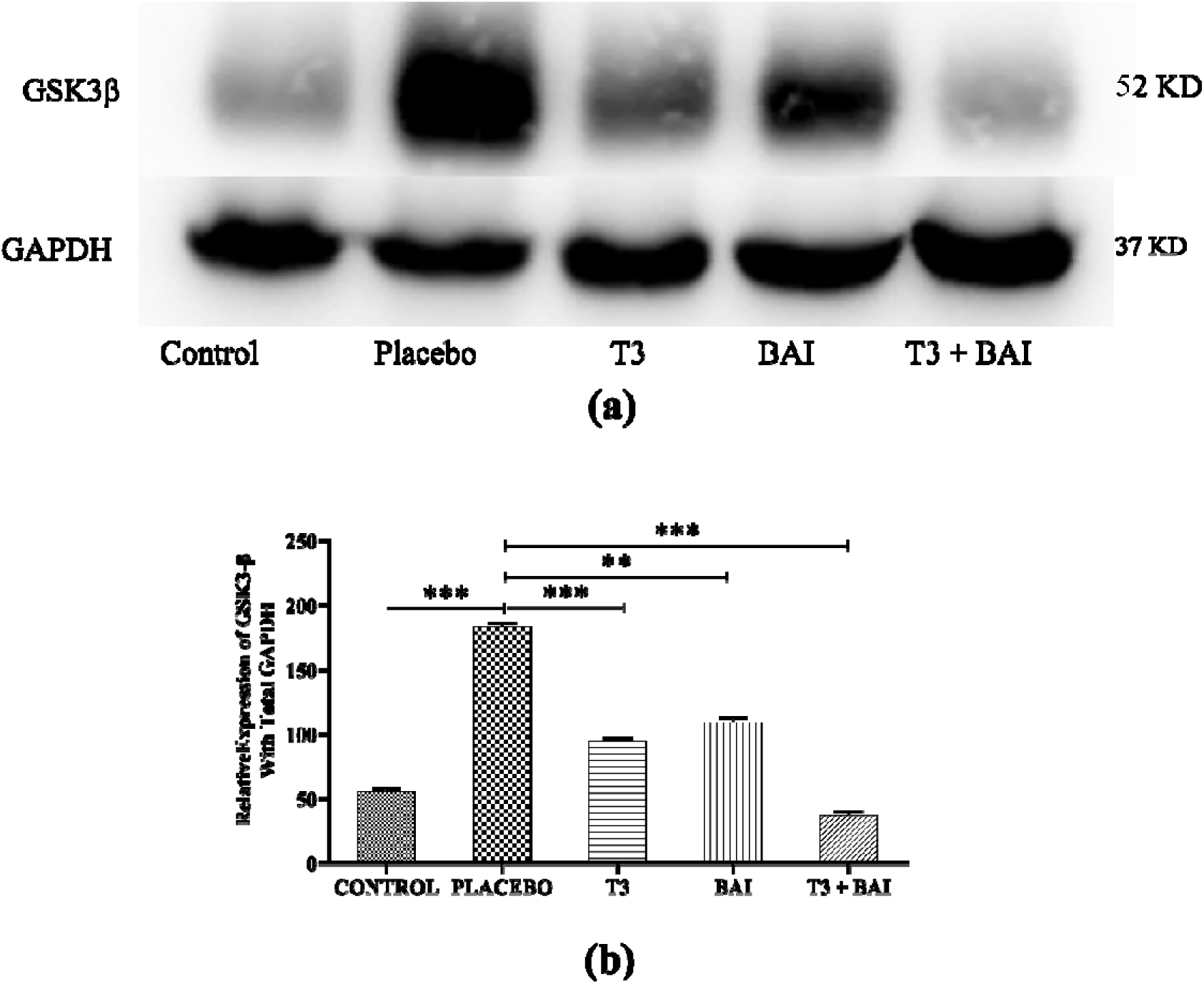
Effects of T3, BAI, and the combination (T3 + BAI) on GSK-3β protein expression. T3 and BAI successfully reduced GSK-3β protein expression, with a greater reduction observed in the combined (T3 + BAI) treatment group. **(a)** and **(b)** Western blot analysis of GSK-3β and densitometric analysis in CKD mice. The data are expressed as the means□±□SEMs from 3 independent experiments. ⍰⍰ and ⍰⍰⍰ indicate significance at p ≤ 0.01 and p ≤ 0.001, respectively.

### Combined treatment (T3 + BAI) suppresses the canonical Wnt/**β**-catenin pathway in CKD animals

Understanding the crosstalk of the evolutionarily conserved Wnt/β-catenin pathway with other signaling pathways is crucial for understanding kidney disease progression and elucidating the therapeutic potential of treatment protocols [48]. To determine the effects of our treatments on the canonical Wnt/β-catenin pathway, the mRNA expression of Wnt ligands such as Wnt1, 3, 8A, 8B, and 10A and the protein expression of β-catenin and casein kinase 1 (CK-1) were measured. We found that the mRNA expression of all Wnt ligands increased significantly in the placebo-treated mice compared with the control mice **(Fig. 6a).** However, treatment with BAI alone or in combination (T3 + BAI) repressed the expression of these mRNAs compared with that in the placebo group **(Fig. 6b).** In contrast, no significant (p<0.99) changes were observed in the group that received only the T3 treatment.

**Fig. 6.**
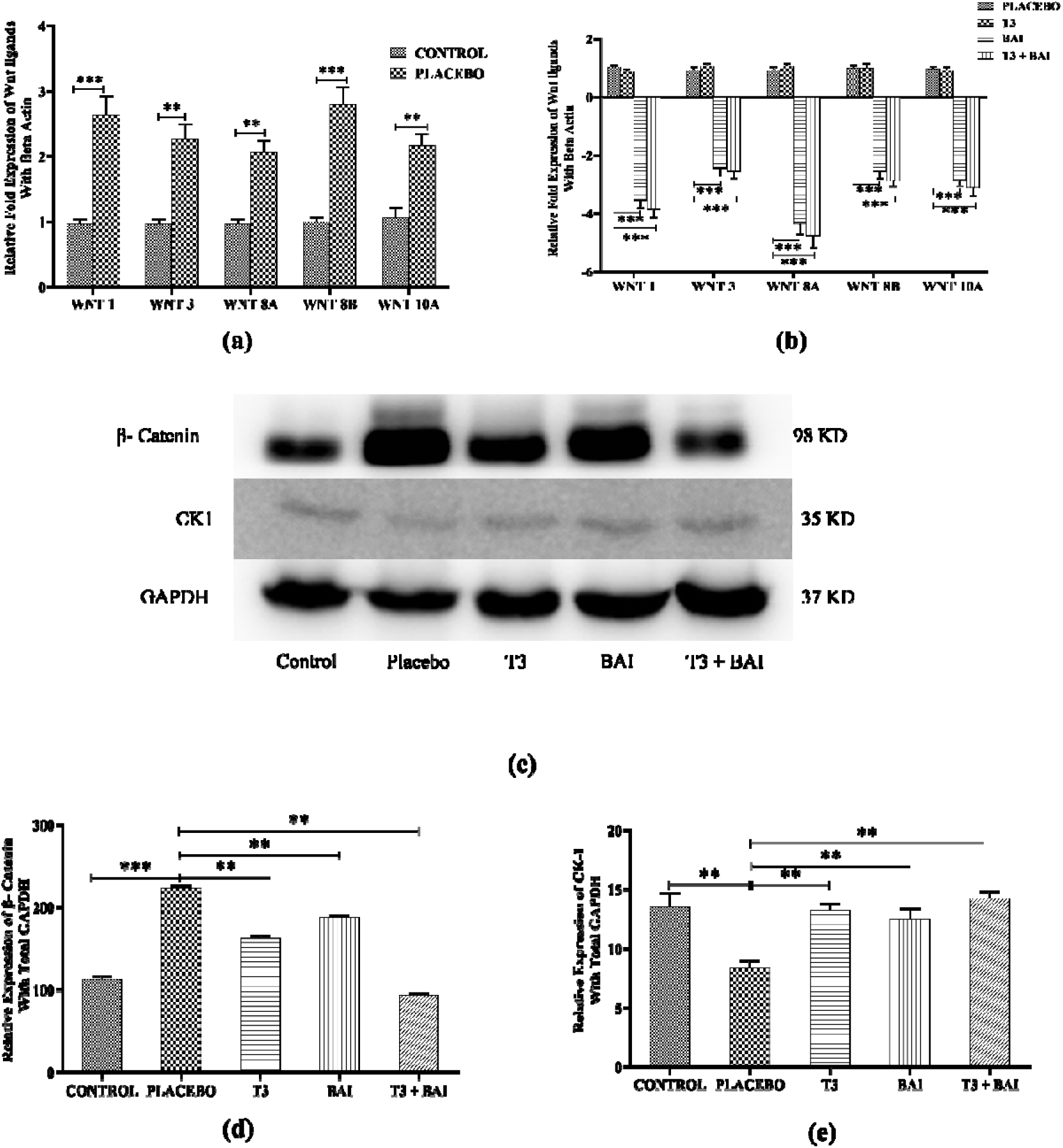
Effects of T3, BAI, and the combined treatment (T3 + BAI) on the Wnt-β-catenin pathway in CKD kidneys. CKD mice presented increased Wingless-related integration site (Wnt) ligand mRNA expression with increased β-catenin protein expression. The downregulation of the expression of all the mRNAs and β-catenin proteins was observed in both the T3 and BAI treatment groups, with the maximum reducing effect observed in the combined treatment group. **(a)** and **(b)** mRNA expression of Wnt ligands in different groups. **(c)** Western blot analysis of β-catenin and creatine kinase 1 (CK-1) expression; **(d)** and **(e)** Densitometric analysis of β-catenin and CK-1 protein expression, respectively. The data are expressed as the means□±□SEMs from 3 independent experiments. ⍰⍰ and ⍰⍰⍰ indicate significance at p ≤ 0.01 and p ≤ 0.001, respectively.

At the level of β-catenin protein expression, we observed a significant increase of 97.9% (p<0.001), and CK-1 expression was 38.5% (p<0.01) lower in the placebo group than in the control group. In contrast, after the T3 and BAI treatments, the protein expression of β-catenin decreased significantly [T3: 27.6% (p<0.01); BAI: 17.8% (p<0.01)], and CK-1 [T3: 58.5% (p<0.01); BAI: 50.4% (p<0.01)] increased significantly toward a normal level. Moreover, the combined treatment group exhibited a maximum decrease in β-catenin [58.3% (p<0.01)] with increasing CK-1 [71% (p<0.01)] protein expression compared with the placebo group. **(Fig. 6c, d, e).**

### Treatments can modulate kidney TGF-**β** expression

Regarding the genesis of CKD, the role of TGF-β as an essential mediator of the whole process has been recognized recently [49]. In the present study, we found that the expression of TGF-β mRNA increased significantly, by 4.1-fold (p<0.001), in the placebo group compared with the control group **(Fig. 7a).** Upon treatment with either T3 and/or BAI, a significant reduction in TGF-β mRNA expression of 2.3-fold (p<0.05) and 2.6-fold (p<0.05) was observed compared with that in the placebo group. In comparison, the combined (T3 + BAI) group presented a 3.56-fold greater reduction in TGF-β mRNA expression (p<0.05) than the placebo group did **(Fig. 7b).**

**Fig. 7.**
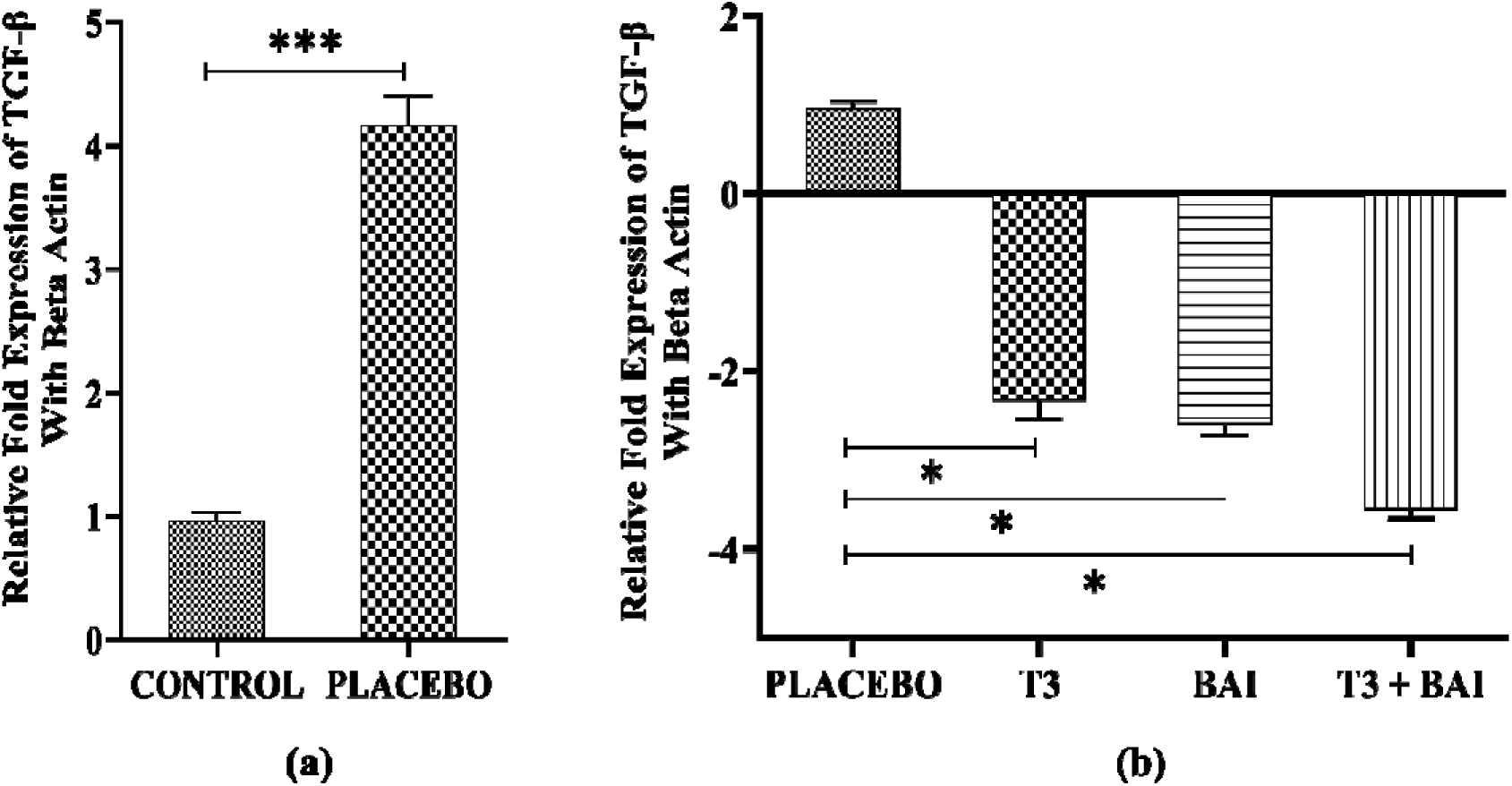
Effects of T3, BAI, or their combination (T3 + BAI) on TGF-β mRNA expression. The abnormal increase in transforming growth factor-β (TGF-β) mRNA expression was reduced by T3 and BAI treatment, with a greater reduction in the combined treatment. The data are expressed as the means□±□SEMs from 3 independent experiments. ⍰ and ⍰⍰⍰ indicate significance at p ≤ 0.05 and p ≤ 0.001, respectively.

### Protective effect against CKD-induced inflammation

A proinflammatory state is also linked to chronic kidney disease, and persistent low-grade inflammation is increasingly recognized as a defining feature of CKD [50,51]. Increased production and decreased clearance of proinflammatory cytokines contribute to the chronic inflammatory status in CKD [52,53]. Thus, we analyzed the expression of cytokines and inflammatory immune cell accumulation in the kidneys to assess the intensity of renal inflammation.

The protein expression levels of inflammatory kidney markers, such as nuclear factor-κB (NF-κB) and interleukin 6 (IL-6), were significantly greater in the placebo-treated mice than in the control mice. This includes increases in NF-κB by 2.7-fold (p≤0.001) and in IL-6 by 3-fold (p≤0.001). Moreover, the T3 and BAI groups presented significantly lower levels of inflammatory markers than did the placebo group. Moreover, compared with the placebo treatment, the individual T3 and BAI treatments significantly reduced N-FκB expression by 2.2-fold (p≤0.01) and 1.8-fold (p≤0.01), respectively. Similarly, compared with the placebo, T3 and BAI significantly reduced IL-6 expression by 1.5-fold (p≤0.01) and 1.65-fold (p≤0.01), respectively. The maximum reduction in inflammatory markers was observed in the combined treatment group, in which NF-κB expression was reduced by 4.3-fold (p≤0.001) and IL-6 expression was reduced by 3-fold (p≤0.001) compared with those in the placebo group **(Fig. 8a, b, c).** These findings provide evidence that combined T3 + BAI could alter the increase in inflammatory cytokines during CKD, thereby protecting the kidneys from further deterioration.

**Fig. 8.**
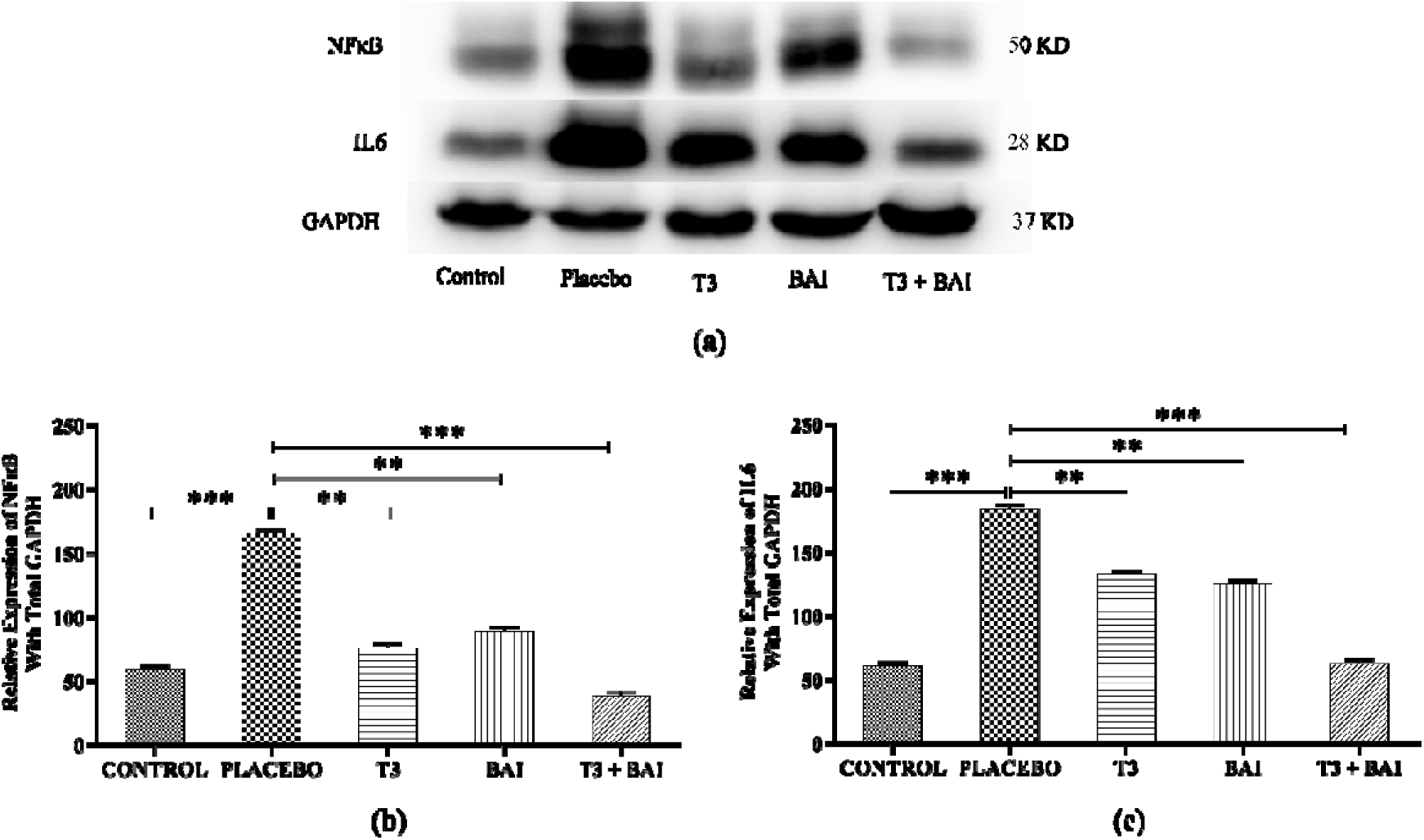
Effects of T3, BAI, and the combined treatment (T3 + BAI) on proinflammatory marker levels in CKD kidneys. The protein expression of proinflammatory markers such as nuclear factor kappa B (NFκB) and interleukin 6 (IL-6) is significantly increased in CKD kidneys. In terms of kidney protection, T3 and BAI reduced both NFκB and IL-6 expression, with the greatest reduction observed in the combined treatment group. **(a)** Western blot analysis of NFκB and IL-6; **(b)** and **(c)** Densitometric analysis of NFκB and IL-6 protein expression, respectively. The data are expressed as the means□±□SEMs from 3 independent experiments. ⍰⍰ and ⍰⍰⍰ indicate significance at p ≤ 0.01 and p ≤ 0.001, respectively.

### T3 but not BAI could protect CKD animals from anemia

A decrease in one or more important red blood cell parameters, such as the hematocrit, hemoglobin concentration, or red blood cell count, is referred to as anemia. Since anemia is a frequent side effect of CKD, it is unquestionably an independent risk factor for worse outcomes in CKD patients. Anemia is less common in early kidney disease, and it often worsens as kidney disease progresses and when more kidney function is lost [54,55]. Hence, we were interested in determining the anemia status in CKD mice and the effect of treatment on anemia status. The hematocrit percentage and hemoglobin concentration were significantly lower in the placebo group than in the control group by 27.4% (p≤0.05) and 23.3% (p≤0.05), respectively. Compared with those in the placebo group, the hematocrit percentages in the T3- and combined (T3 + BAI)-treated groups were 17.08% (p≤0.05) and 15.7% (p≤0.05), respectively. Similarly, the Hb concentration increased by 14.7% (p≤0.05) and 14.7% (p≤0.05), respectively. In contrast, no significant changes were observed in the BAI-treated group (p≤0.48). We can conclude that only T3 could protect animals from CKD-induced anemia (**Table 4)**.

**Table 4.** Serum parameters for anemia.

### T3 but not BAI modulates mineral bone disorders in CKD

CKD-induced mineral bone disorders include abnormalities in bone and mineral composition [56]. CKD patients develop an inadequate 1,25-dihydroxyvitamin D level, predisposing them to hyperphosphatemia. Furthermore, both processes together cause a reduced serum calcium level [5]. As a result of bone wear and tear, the serum alkaline phosphatase (ALP) level is increased in CKD patients [57–59]. Hence, serum phosphate, calcium, vitamin D3, and alkaline phosphatase (ALP) activities were estimated to investigate the potential of treatments for modulating mineral and bone disorders. Our results revealed a significant increase in the serum phosphate concentration and a decrease in the calcium concentration in the placebo-treated mice compared with those in the control mice (**Table 5)**.

**Table 5.**
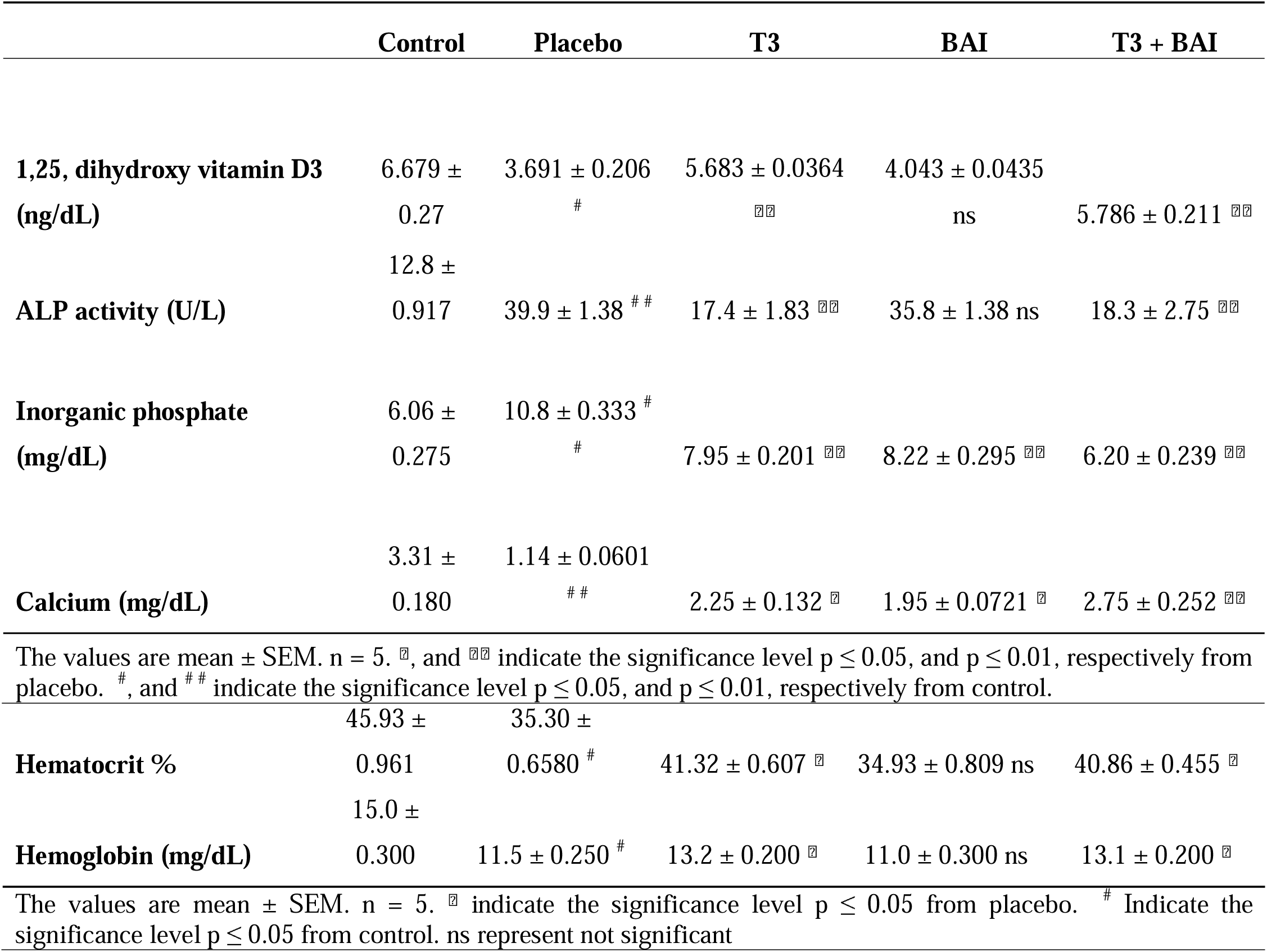
Effects on the serum parameters for minerals and Bone disorders.

Moreover, these changes in Ca and Pi levels during CKD were reversed toward their normal range by the T3 and BAI treatments, with maximum reversal caused by the combined treatment. Similarly, an increase in the mRNA expression of the phosphate channel NPT2a, followed by a decrease in the expression of the calcium channel TRPV5, was detected in CKD animals. However, the combined treatment had a much greater positive effect on the altered changes caused by CKD, reversing the abnormality through decreased NPT2a and increased TRPV5 mRNA expression **(Figure 9)**. Compared with those in control mice, a significant decrease in serum vitamin D3 levels and an increase in alkaline phosphatase (ALP) activity were also observed in placebo-treated mice, indicating bone wear and tear. However, these abnormal changes were counteracted by T3 and combined (T3 + BAI) treatment. No significant differences were observed in the BAI-treated group (p≤0.34) (**Table 5)**.

**Fig. 9.**
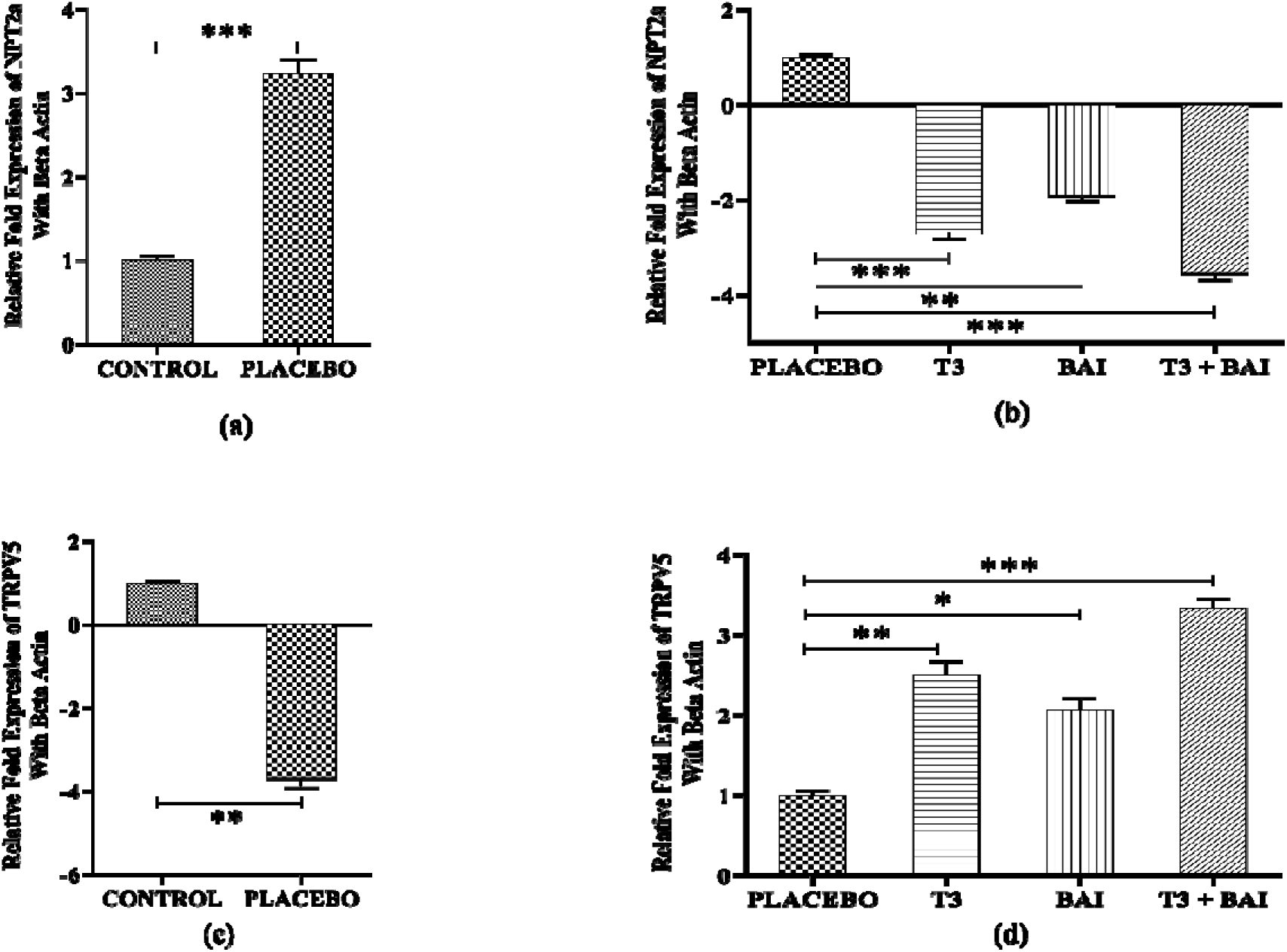
Effects of T3, BAI, and the combined treatment (T3 + BAI) on the mRNA expression of phosphate and calcium transporters. T3 and BAI treatment significantly protected the kidney and altered mRNA expression, with a more pronounced effect in the combined treatment group. **(a)** and **(b)** mRNA expression of NPT2a in different groups and treatments. **(c)** and **(d)** represent the mRNA expression of TRPV5. The data are expressed as the means□±□SEMs from 3 independent experiments. IZ, IZIZ, and IZIZI Z indicate significance at p ≤ 0.05, p ≤ 0.01, and p ≤ 0.001, respectively.

### Protective effect on CKD-associated cardiovascular risk

End-stage renal disease (ESRD) is widely known to increase cardiovascular risk, and it is estimated that dialysis patients have cardiovascular death rates that are ten to one hundred times greater than those in the general population [60]. A heightened proinflammatory microenvironment and other modifying factors (dyslipidemia) during CKD predispose individuals to cardiovascular events **(Supplementary Result S3)**. The atherogenic index (AI) and coronary risk index (CRI), which are calculated on the basis of serum triglyceride, total cholesterol (TC) and high-density lipoprotein cholesterol (HDL-C) levels (**Supplementary Table 1)**, were significantly greater (p≤0.001) in the placebo group than in the control group **(Figure 10a, b)**. The risk levels pertaining to the AI and CRI were lower in the T3- and BAI-treated animals than in the placebo**-**treated animals. Compared with either of the individual treatments, the combined treatment was better at lowering the AI and CRI predispositions.

**Figure 10.**
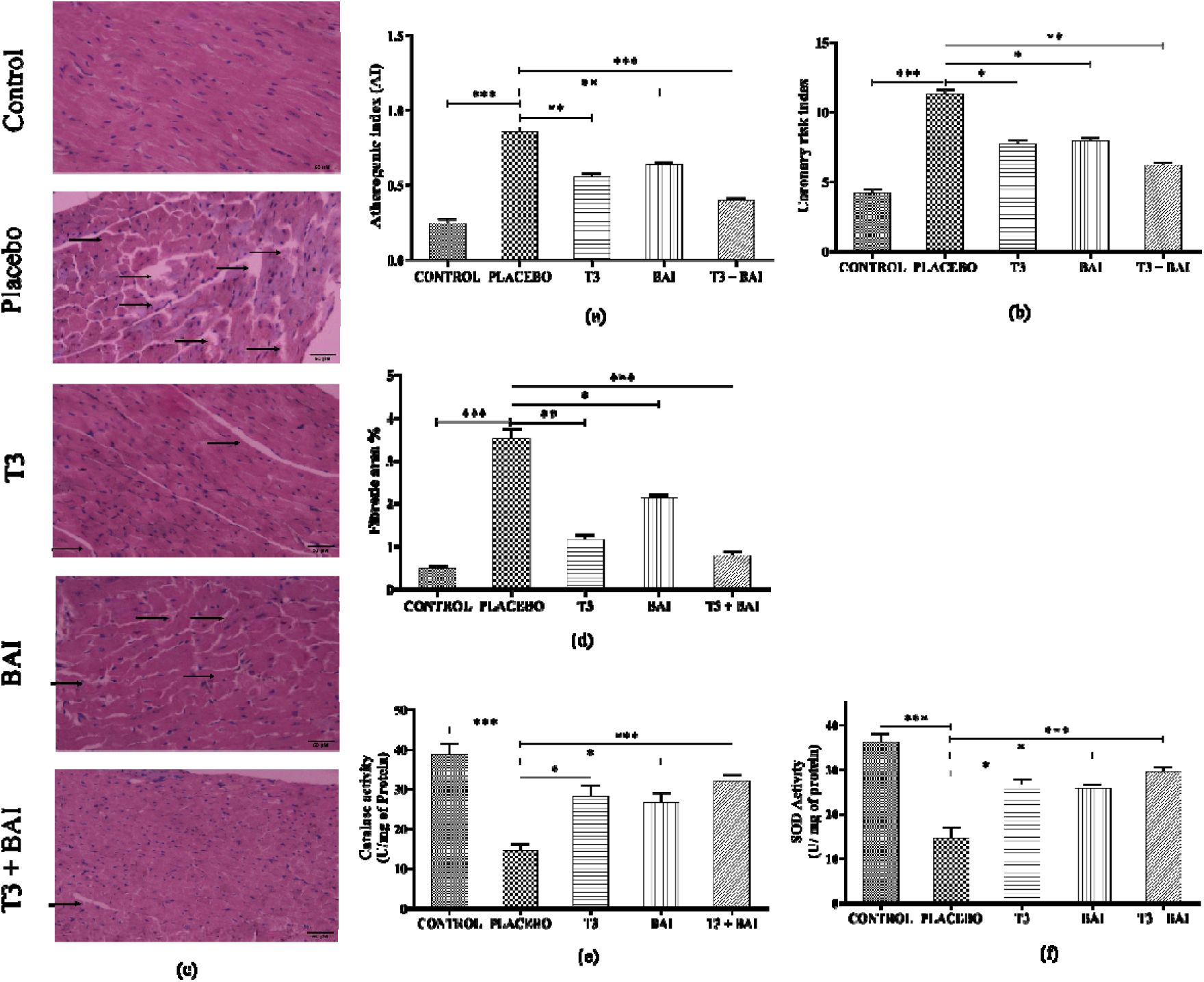
Protective effect against CKD-induced cardiovascular disorder (CVD). T3 and BAI had protective effects against CKD-induced cardiovascular damage, with more pronounced effects observed with T3 treatment. However, maximum protection was observed in the combined (T3 + BAI) treatment. The combined treatment also protected against oxidative damage in heart. **(a)** and **(b)**, The calculated atherogenic index (AI) and coronary risk index (CRI) increased significantly in the CKD placebo group. All the risks for the heart were reduced upon T3 and BAI treatment, with the maximum protective effect observed with the combined dose. **(c)** Histopathological analysis of kidney sections with hematoxylin and eosin (H&E) staining indicating myocardial damage in CKD patients. **(d)** Represents the calculated fibrotic area in the heart section via ImageJ **(e)** and **(f)** represents catalase and SOD activity in heart tissue respectively. ⍰, ⍰⍰, and ⍰⍰⍰ indicate significance at p ≤ 0.05, p ≤ 0.01, and p ≤ 0.001, respectively.

Like kidney tissue, heart tissue was also checked for damage via H&E staining of thin sections. We found that in the placebo group, there was a significant increase in fibrosis and inflammatory infiltration compared with those in the control group. Surprisingly, individual T3 and BAI treatments reduced fibrosis, and T3 had a greater antifibrotic effect than BAI did. Moreover, the combined treatment (T3 + BAI) produced a significantly more pronounced antifibrotic effect than either of the individual treatments did **(Figure 10c, d)**. Like those in the kidney, a similar trend of antioxidant enzyme activities in the heart tissue was observed, wherein SOD and catalase activities were significantly reduced in the placebo-treated mice (p≤0.001). In contrast, T3 and BAI alleviated the activity of both enzymes, and compared with the placebo, the combined treatment (T3 + BAI) resulted in the highest activity of both enzymes.

## Discussion

In the present study, we aimed to decipher the protective effect of combined T3 and BAI treatment on adenine-induced CKD in old mice. As per our previous findings [17,61], we intend to bidirectionally target CKD using the thyroid hormones T3 and BAI. T3 was used with the intention of maintaining a normal HPT axis in CKD, whereas BAI was used to counteract the exaggerated upregulation of the Wnt/β-catenin pathway, as previously reported. Interestingly, we found that both treatments could enhance Klotho via different targets. Furthermore, as CKD predisposes individuals to other complications, such as anemia, dyslipidemia, cardiovascular disease, and mineral bone disorders, we intend to evaluate the therapeutic potential of individual (either T3 or BAI) versus combined (T3 + BAI) treatment for these complications.

Serum and urine biomarkers, including creatinine, urea, BUN, and albumin, are used as standard clinical indices to assess kidney disorders leading to CKD [51,62–64]. The adenine oral dose significantly impaired kidney function, as evidenced by increased serum creatinine, urea, BUN, and urine albumin levels, followed by decreased urinary excretion of creatine and urea. In this study, when CKD mice were treated with T3, BAI, or their combination (T3 + BAI), all the abnormal serum parameters reverted toward their normal range, indicating that there were improvements in kidney function after the treatments.

Research indicates that CKD significantly impacts the HPT axis, impairing thyroid hormone metabolism [5,65,66]. Subclinical hypothyroidism prevalence rises with decreasing GFR and advancing age, often accompanied by lower serum T3 and T4 levels and higher TSH levels [7]. Our findings align with these reports, showing reduced T3 and elevated TSH in CKD mice (**Supplementary Results S1)**.

To address HPT axis dysregulation, exogenous T3 was administered at a previously reported dose [35,36]. After 12 hours of treatment, TSH levels significantly decreased in the T3 and T3 + BAI groups, while no notable changes occurred in BAI-only animals due to T3’s biological half-life. Further studies are needed to refine the dosage and timing of thyroid hormone therapy, given the study’s limitations in pharmacokinetic analyses in CKD models.

Glomerulosclerosis, interstitial fibrosis, and tubular atrophy are symptoms of renal fibrosis resulting from the inability of kidney tissue to be repaired after long-term, chronic damage [67]. Renal cells are replaced by ECM in the glomeruli and interstitium as CKD progresses, which is accompanied by the loss of renal cells [68,69]. Morphological analysis of CKD kidneys using H&E and Masson’s trichrome staining revealed increased inflammatory infiltration, protein casts, and ECM deposition following adenine treatment. However, T3 and BAI combination therapy significantly reduced fibrosis and ECM deposition, demonstrating a protective effect against further deterioration in aged CKD mice.

Klotho is primarily expressed on the cell surface of DCT and PCT in the kidney and is maintained at normal levels under physiological conditions [70–74]. However, CKD leads to reduced Klotho levels, evident in both nephritic dysfunction and animal models. CKD decreases mRNA expression of M-Kl and Sp-Kl and reduces α-Kl and S-Kl protein levels [75,76].

T3 and BAI treatments restored Klotho mRNA and protein expression toward normal, with the combination treatment showing the greatest effect. BAI likely increases Klotho by inhibiting the Wnt/β-catenin pathway, while T3 directly activates Klotho, as suggested by previous findings.

The standard progression route for CKDs, including renal tubulointerstitial fibrosis, often leads to ESRD. Recent findings highlight GSK3β’s role in renal tubular cell profibrogenic plasticity during CKD progression, particularly through its interaction with pathways like TGF-β1 signaling [46]. Elevated GSK-3β expression in glomeruli and renal tubular cells is linked to kidney diseases like proteinuric glomerulopathies and podocyte damage [77–79]. Determining changes in GSK3β expression after treatment is essential. CKD kidneys show elevated GSK-3β levels, indicating tubular inflammation and damage. While individual treatments with T3 and BAI downregulated GSK-3β, they did not significantly reduce fibrosis. In contrast, combined therapy markedly reduced GSK-3β expression, likely by stabilizing Klotho expression. T3 enhances Klotho levels, while BAI reduces its transcriptional inhibition by β-catenin, as Klotho downregulates GSK-3β expression [80].

The canonical Wnt/β-catenin pathway, an evolutionarily conserved signaling pathway, controls the pathogenesis of many diseases. Overexpression of this pathway has been observed in different types of kidney injury [81–84]. Sustained activation of the canonical Wnt/β-catenin pathway drives acute kidney injury (AKI) to CKD progression [85]. Adenine-fed animals displayed enhanced activation of the Wnt/β-catenin pathway, as evidenced by increased mRNA expression of various Wnt ligands, reduced CK-1 protein levels, and elevated β-catenin protein production in the kidney. The natural compound BAI effectively suppressed the elevated mRNA expression of all Wnt ligands in CKD mice. Moreover, T3 and BAI effectively restored β-catenin and CK-1 expression levels in CKD mice, with the combination therapy further promoting β-catenin degradation. We propose that T3’s inhibitory effect on the Wnt pathway may be mediated through its ability to upregulate Klotho expression, as Klotho can inhibit the Wnt/β-catenin pathway without directly affecting Wnt ligand mRNA expression. (Add t3 Klotho paper) [86–88]. Similarly, since the highest levels of Klotho expression were observed with the combined T3 and BAI treatment, we speculate that BAI’s inhibition of the Wnt/β-catenin pathway may prevent this major upregulated pathway, observed in CKD, from suppressing Klotho expression. This mechanism likely ensures the sustained upregulation of Klotho protein in the kidney, contributing to its pronounced beneficial effects.

TGF-β is a key profibrotic regulator in CKD, promoting ECM accumulation, preventing its breakdown, and activating myofibroblasts [89–92]. Therapeutic strategies targeting TGF-β in experimental and clinical CKD models have shown reduced renal fibrosis and injury [93,94]. Studies in renal epithelial cells reveal that Klotho inhibition increases TGF-β1 expression, while TGF-β1 suppresses Klotho, indicating that reduced Klotho promotes TGF-β1 activity, creating a cycle contributing to renal fibrosis in CKD [95]. In this study, adenine-induced CKD mice showed increased TGF-β mRNA expression, linked to activation of the Wnt/β-catenin pathway and reduced Klotho expression. However, the combination treatment group exhibited a significant reduction in TGF-β levels. These findings align with previous reports [96], suggesting that Klotho inhibits TGF-β signaling, offering potential benefits in suppressing renal fibrosis.

Fibrosis in CKD is closely associated with increased inflammation, characterized by elevated levels of inflammatory cytokines such as NF-κB, TNF-α, and interleukins in adenine-induced CKD mice. Klotho, as an anti-inflammatory regulator, can mitigate NF-κB activity by reducing proinflammatory gene transcription [97]. Consistently, our study observed a marked increase in NF-κB and IL-6 protein expression in the kidneys of CKD mice. We hypothesize that our dual treatment strategy may counteract inflammation by enhancing Klotho expression. Notably, the anti-inflammatory effects of T3 and BAI were most pronounced in the combination therapy (T3 + BAI) group, potentially attributable to the significant upregulation of Klotho expression induced by the combined treatment.

CKD is marked by oxidative stress due to reduced antioxidant activity and excessive ROS production [98], with a strong link between ROS generation and Klotho levels. Klotho mitigates oxidative stress by downregulating insulin/IGF-1/PI3K signaling, activating FoxO, and inducing MnSOD formation [99]. Additionally, free radicals (e.g., H2O2, O2•–) are generated during hypoxanthine/xanthine conversion to uric acid via xanthine oxidase (XO) [100–102]. Adenine-induced overproduction of hypoxanthine elevates XO expression, causing oxidative damage in CKD kidneys [103]. To evaluate the kidney’s antioxidant defense, we measured catalase and SOD activity. As anticipated, adenine-induced CKD kidneys showed reduced expression of these antioxidant enzymes. All treatment groups effectively restored their altered activity levels. (**Supplementary Results S2)**.

CKD is associated with various complications, including anemia, mineral and bone disorders, dyslipidemia, and cardiovascular conditions. We investigated whether our dual treatment offers protection solely against CKD or also mitigates these associated complications. Notably, the combined treatment demonstrated protective effects against all the aforementioned CKD-related complications.

In the case of CKD-induced anemia, which arises from mechanisms such as reduced erythropoietin synthesis, hypothyroidism, and shortened red blood cell lifespan [5,104], the protective effect was mediated by T3 but not by BAI. This can be attributed to T3’s well-established role in stimulating erythropoiesis [105–107].

To examine its effect on mineral and bone disorders, we measured phosphate (Pi) and calcium (Ca²□) levels, commonly altered in CKD (increased Pi and decreased Ca²□ levels) [15]. Since Klotho regulates phosphate and calcium homeostasis, our combined treatment normalized these levels via the actions of both T3 and BAI. This was further supported by increased mRNA expression of kidney phosphate and calcium channels regulated by Klotho. Mineral and bone disorders in CKD are also driven by elevated alkaline phosphatase (ALP) activity and reduced vitamin D3 levelsv [108].. These abnormalities were corrected by T3, highlighting its additional protective role. Thus, T3 and BAI function through distinct mechanisms to address CKD-induced anemia and mineral and bone disorders.

CKD is a systemic condition often accompanied by dyslipidemia. As renal function declines, hyperlipidemia becomes more common, with elevated LDL cholesterol and triglyceride levels correlating with the severity of impairmen [5]. Furthermore, dyslipidemia and elevated serum TSH levels are independent risk factors for cardiovascular diseases [109,110].

The bidirectional treatment effectively reversed the altered lipid profile and protected against dyslipidemia (**Supplementary Results S3)**. To evaluate its protective effect on CVD, we initially assessed coronary disease risk using lipid profile markers by calculating the Atherogenic Index (AI) and Cardiac Risk Index (CRI). CKD mice showed a higher CVD risk, which decreased with T3 and BAI treatments, with the combined treatment providing the greatest reduction.

Given the elevated CVD risk in CKD mice, we examined heart tissue for fibrosis using H&E staining. Notably, CKD hearts displayed significant fibrosis. Similar to findings in the kidney, combined treatment offered superior protection in the heart compared to individual therapies, as evidenced by reduced fibrotic areas, highlighting its protective role against CVD.

Increased oxidative stress in the kidney prompted an evaluation of antioxidant enzyme activity in heart tissue. Both catalase and SOD activities were diminished in CKD mice but improved with treatment, demonstrating the protective effects of Klotho in cardiac tissue as well.

## Conclusion

This study identified a novel therapeutic approach for CKD treatment using the endogenous hormone T3 and the natural compound BAI. Both agents stabilize Klotho protein in CKD kidneys through distinct pathways. The reduced Klotho expression in CKD results from decreased T3 levels and activation of the Wnt/β-catenin pathway. This combined therapy shows potential for greater efficacy, particularly in older patients with subclinical hypothyroidism.

## Ethics approval

Approved by the Institutional Animal Ethics Committee (IAEC) of Pondicherry University, India (IAEC approval ref. No: PU/CAHF/23rd IAEC/2019/03, dated 15.04.2019

## Acknowledgments

The authors acknowledge the Department of Biochemistry and Molecular Biology, Pondicherry University, India, and the Department of Science & Technology “Fund for Improvement of S&T Infrastructure”, (DST-FIST), Science and Engineering Research Board (SERB), Government of India, for financial and overall support.

## Credit authorship contribution statement

**Saswat Kumar Mohanty**: Conceptualization, methodology, investigation, formal analysis, writing - original draft, project administration

**Vikas kumar Sahu, Bhanu Pratap Singh:** Helped SKM in conducting investigations as a part of their M.Sc. dissertation.

**Kitlangki Suchiang**: Supervision, conceptualization, validation, formal analysis, review and editing, funding acquisition

## Data availability

The datasets generated and/or analyzed in this study are available from the corresponding author Kitlangki Suchiang upon reasonable request.

## Abbreviation

T3: Triiodothyronine
BAI: Baicalein
M-Kl: Membrane Klotho
S-Kl: Soluble Klotho
Sp-Kl: Spliced Klotho
α-Kl: Alpha Klotho
Wnt: Wingless-related integration site
AI: Atherogenic index
BUN: Blood urea nitrogen
CKD: Chronic kidney disease
CVD: Cardiovascular disease
GSK-3β: Glycogen synthase kinase 3 beta
TRPC6: Transient receptor potential cation channel
TRPV5: Transient receptor potential cation channel subfamily V member 5
TGF-β: Transforming growth factor-beta
TC: Total cholesterol
TG: Triglyceride
LDL: Low-density lipoprotein
V-LDL: Very low-density lipoprotein

