## Supplementary material for "Bidirectional Upregulation of Klotho by Triiodothyronine and Baicalein: Mitigating Chronic Kidney Disease and Associated Complications in Aged BALB/c Mice": Suplimentary Information

**Supplementary information**

**Authors**

**Saswat Kumar Mohanty*****^1^, Vikas Kumar Sahu^1^, Bhanu Pratap Singh^1^, Kitlangki Suchiang^2^,**

^1^Department of Biochemistry and Molecular Biology

Pondicherry University, Pondicherry – 605 014, India

^2^Department of Biochemistry, North Eastern Hill University

*****

***** orcid: [orcid.org/0000-0002-6155-3894](https://orcid.org/0000-0002-6155-3894)

**Supplementary Results**

**Supplementary Results S1. CKD changes the serum thyroid hormones profile**

Subclinical hypothyroidism or low T3 syndrome is the most common laboratory finding in CKD patients, as reported elsewhere ^1,2^. Our study found that the serum T3 concentration significantly decreased by 1.98-fold (p<0.01) in the placebo group compared to the control group. Treatment with T3 alone and in combination with BAI significantly increased serum T3 concentration by 4.14 and 4.11-fold (p<0.001), respectively, from the placebo group after 12 hrs of administration. Similarly, the TSH concentration increased significantly in the placebo group from the control group by 1.38-fold (p<0.01). On treatments with either T3 and combined (T3 + BAI), a decline of 1.28 and 1.30-fold (p<0.001), respectively, was observed in comparison to the placebo group. As expected, no significant changes were observed in the serum thyroid hormone profile of BAI-treated mice (p<0.57). **(Supplementary Fig. S1)**

**Supplementary Results S2. T3 + BAI protects the kidney from oxidative damage**

Impaired antioxidant enzyme activity has been observed in CKD, which leads to kidney damage ^3^. We found that the catalase and SOD enzyme activity reduced significantly in the placebo group. The percentage of decrease was 60.7% (p<0.001) and 52.3% (p<0.01) receptively. Individual T3 and BAI-treated groups altered the changes in enzyme activity by adenine treatment toward normal value. Catalase activity increased by 78.5% (p<0.05) and 98.6% (p<0.05), whereas SOD activity increased by 56.2% (p<0.05) and 62.7% (p<0.05) in T3 and BAI treated groups. In comparison, the combined treatment produced the highest catalase and SOD activity with an increment of 125% (p<0.001) and 87.5% (p<0.01), respectively, in comparison to the placebo group **(Supplementary Fig. S2).**

**Supplementary Results S3. Protective effect against CKD-induced dyslipidaemia**

Dyslipidaemia is common among patients with CKD. When renal function diminishes, the incidence of hyperlipidaemia rises, with the severity of hypertriglyceridemia and LDL cholesterol elevation correlating to the severity of renal impairment. ^4^. Serum total cholesterol (TC) and triglyceride (TG) levels increased significantly in the placebo group compared to the control group. Moreover, an unfavourable dysregulation was observed with LDL, and V-LDL levels increased significantly compared to the declined levels of HDL in the placebo group. These changes were reverted toward the commonly accepted levels after T3 and BAI treatment. However, in all these dysregulations, lipid profile parameters have been normalized maximally in a group that received a combination of T3 and BAI treatment, with values ranging toward their normal range (**Supplementary Table 1.)**.

**Supplementary table:**

| **Table 1.** | | | | | |
| --- | --- | --- | --- | --- | --- |
|  | **Control** | **Placebo** | **T3** | **BAI** | **T3 + BAI** |
| **Total Triglyceride (mg/dL)** | 100.0 ± 2.804 | 210.3 ± 4.673 ^# #^ | 141.1 ± 2.804 ^✱✱✱^ | 155.1 ± 3.738 ^✱✱✱^ | 124.3 ± 2.804 ^✱✱✱^ |
| **Total Cholesterol (mg/dL)** | 163.2 ± 5.263 | 281.6 ± 7.895 ^# #^ | 205.3 ± 5.263 ^✱^ | 226.3 ± 5.263 ^✱✱^ | 186.8 ± 2.632 ^✱✱^ |
| **HDL-C (mg/dL)** | 54.55 ± 2.424 | 27.88 ± 1.212 ^#^ | 40.61 ± 1.818 ^✱^ | 36.97 ± 1.818 ^✱^ | 47.88 ± 1.818 ^✱^ |
| **LDL-C (mg/dL)** | 23.74 ± 0.385 | 136.9 ± 3.598 ^# # #^ | 71.49 ± 0.363 ^✱✱✱^ | 85.79 ± 1.889 ^✱✱✱^ | 50.74 ± 4.411 ^✱✱✱^ |
| **VLDL-C (mg/dL)** | 15.85 ± 0.086 | 45.49 ± 0.107 ^# # #^ | 25.55 ± 0.699 ^✱✱✱^ | 29.64 ± 0.191 ^✱✱✱^ | 21.21 ± 0.412 ^✱✱✱^ |
| The values are mean ± SEM. n = 5. ^✱^, ^✱✱^, and ^✱✱✱^ indicate the significance level p ≤ 0.05, p ≤ 0.01 and p ≤ 0.001, respectively from placebo. ^#^, ^# #^, and ^# # #^ indicate the significance level p ≤ 0.05, p ≤ 0.01 and p ≤ 0.001, respectively from control. | | | | | |

**Supplementary Figures:**

**(a)**

**(b)**

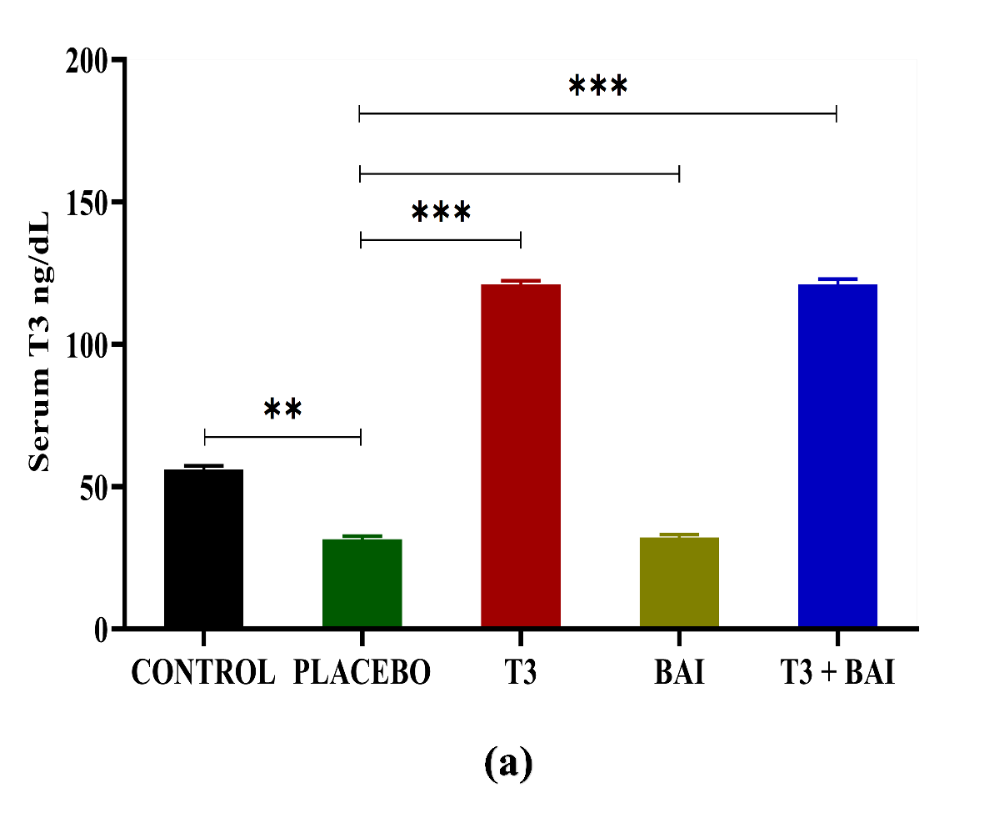

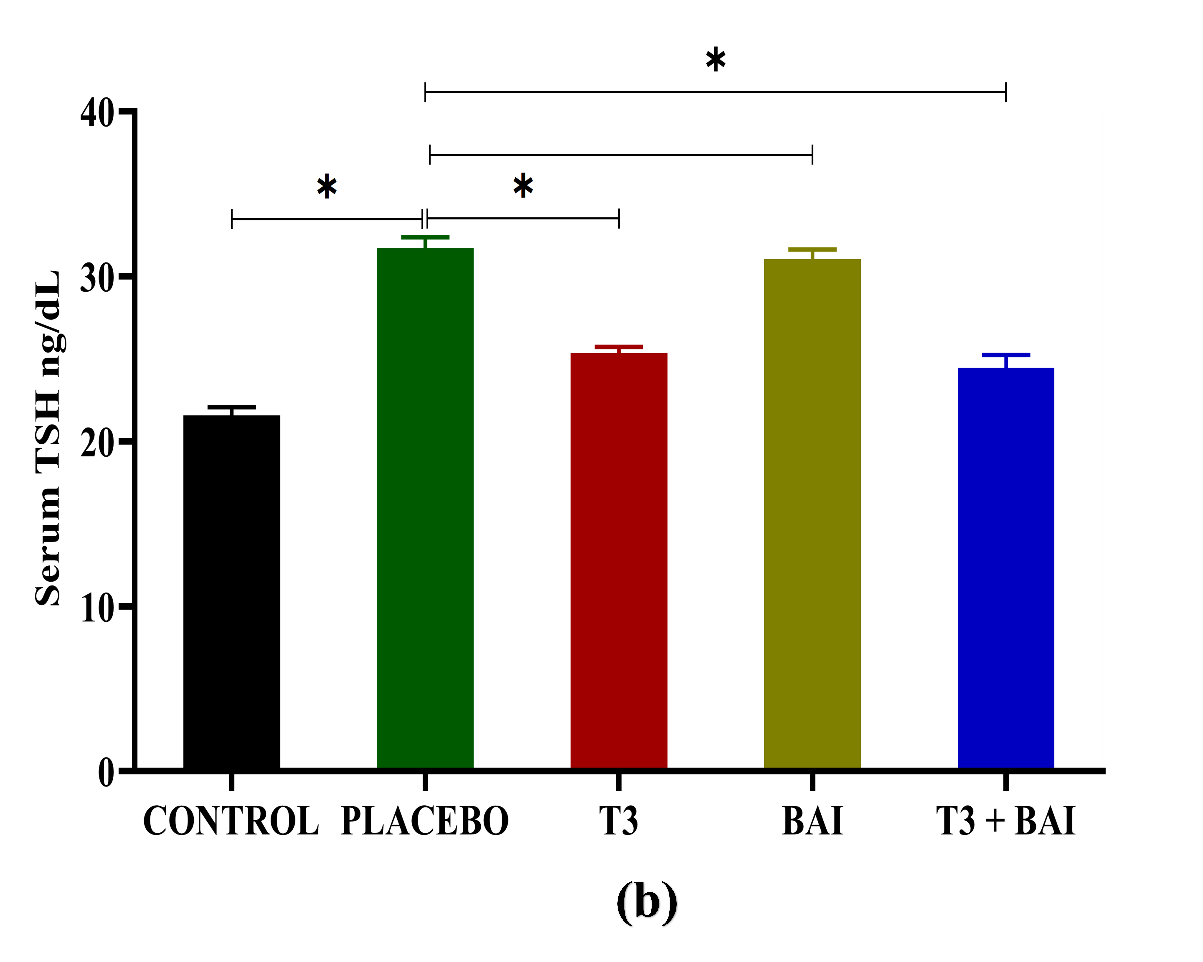

**Fig. S1.**

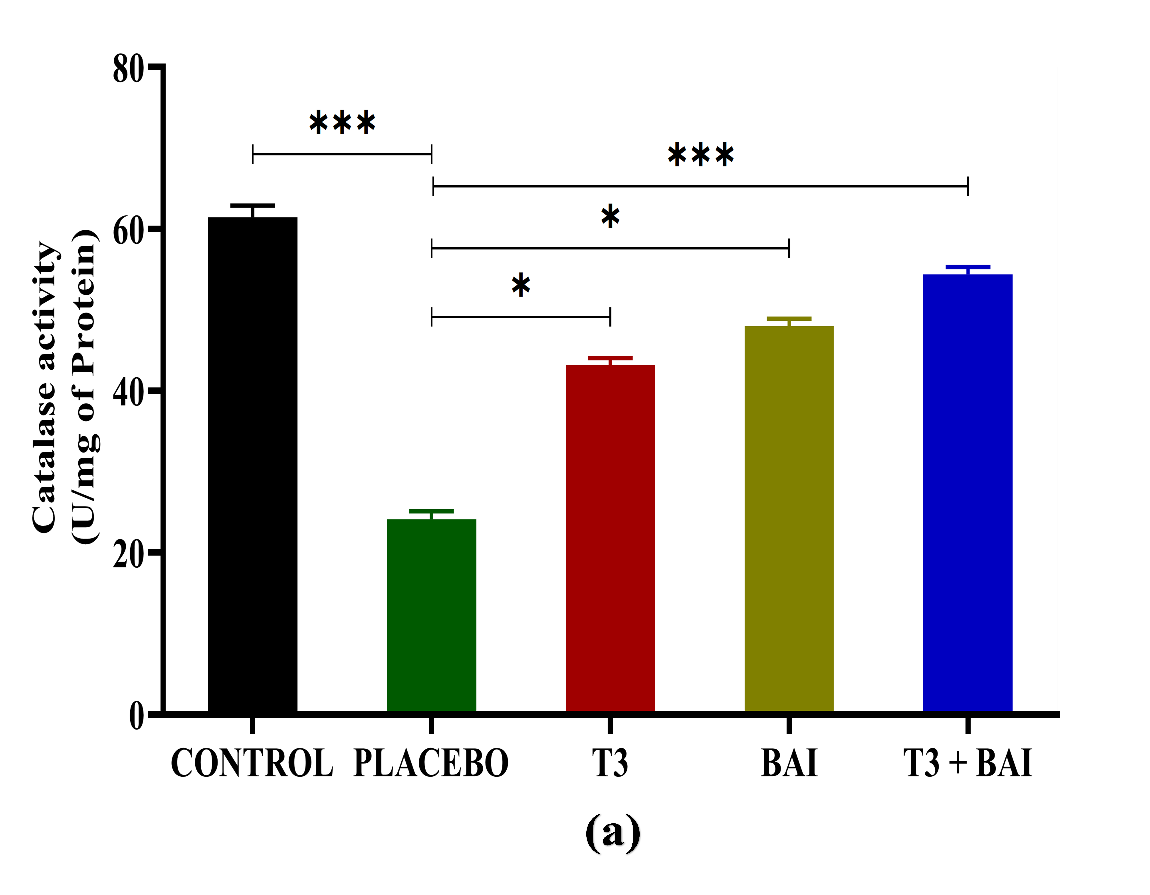

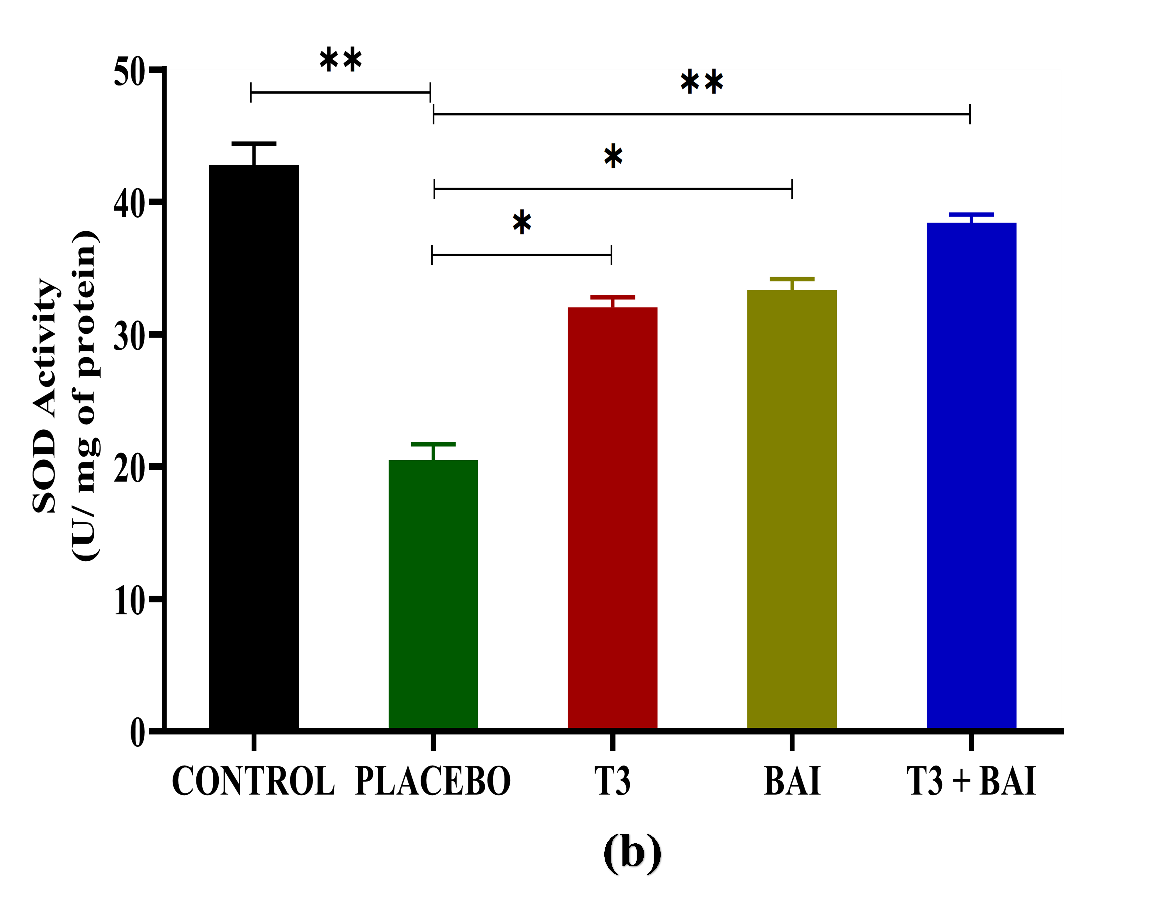

**(b)**

**(a)**

**Fig. S2.**

**Supplementary figure legends:**

**Fig. S1. Effect of T3, BAI, and combined (T3 + BAI) on serum thyroid hormone profile.** Serum T3 concentration was measured by Competitive Elisa and TSH by Sandwich Elisa. T3 concentration decreased, and TSH concentration increased in placebo mice. Reversing effect was observed after T3 treated mice compared to the placebo mice. Data are expressed as means ± SEM from 3 independent experiments. ✱, ✱✱, and ✱✱✱ indicate the significance level at p ≤ 0.05, p ≤ 0.01, and p ≤ 0.001, respectively.

**Fig. S2. Effect of T3, BAI, and combined (T3 + BAI) on kidney antioxidant enzymes.** Activity levels of antioxidant enzymes like catalase (CAT) and superoxide dismutase (SOD) declined in CKD kidneys. On the contrary, T3 and BAI treatment enhanced the activity of both CAT and SOD, with maximum activity observed in the combined treatment. **(a)** CAT activity, **(b)** SOD activity. Data are expressed as means ± SEM from 3 independent experiments. ✱, ✱✱, and ✱✱✱ indicate the significance level at p ≤ 0.05, p ≤ 0.01, and p ≤ 0.001, respectively.

**Supplementary Table legends:**

**Supplementary Table 1.** Effect of T3, BAI, and combined (T3 + BAI) on serum lipid profile parameters
